# Duality Vaccination through Nanodesign of Click-Ligated B-Cell Inhibitory Receptor Bridging

**DOI:** 10.64898/2026.10.28.762652

**Authors:** Gabrielle Admans, Rory A. Hills, Diana Melnyk, Alexander A. Cohen, Claudia L. Driscoll, Mark S. Cragg, Pamela J. Bjorkman, Tiong Kit Tan, Mark R. Howarth

## Abstract

Synthesis of precise and robust molecular architectures is critical to harness the power of the immune system. The dimensions, rigidity and multivalency of nanoparticles contribute to eliciting high levels of antibodies from vaccination. However, antibodies vary hugely in quality: some antibodies are highly effective at neutralizing pathogens, while others provide little or no benefit. Here we establish switchable and modular platforms for protein and nanoparticle assembly that program the target of antibody responses. We introduce Duality vaccination: the simultaneous delivery of activating and inhibitory stimuli for focusing immune responses. To construct activatory immunogens, we employ click biology with spontaneous amidation to functionalize plug-and-display self-assembling porous dodecahedral nanocages. We construct inhibitory immunogens using SpyMask protease-gated bispecific assembly, ligating the antigen to binders that bridge the B cell receptor to immune checkpoint receptors. This dual immunogen architecture tailors the elicited antibodies to down-modulate undesired B cell responses. We establish the effect of temporal and spatial separation of activatory and inhibitory immunogen delivery. We further explore sensitivity to valency, linker flexibility and adjuvant delivery, employing model targets and coronavirus antigens. Dual programmable functionalization identifies key nanodesign features to control *in vivo* outputs, providing logic-gating towards urgent challenges in the creation of protective vaccines.

## Introduction

In the field of molecular recognition, there is still much to learn from the mammalian immune system (Ausserwöger *et al*, 2022). After exposure to micrograms of target antigen, mammals can generate a range of specific nanomolar affinity target binders, showing minimal non-specific binding to the thousands of other molecular targets in the organism (Ausserwöger *et al*, 2022; Graham *et al*, 2019). For viruses there may be only a few target antigens but for bacteria and parasites there may be hundreds (Wood & Wright, 2019). By comparing vaccine immunogens and the mechanism of therapeutic antibodies, it has been established that some antibodies are highly effective in protecting from infection, such as those blocking receptor binding of primary viral receptors for cell attachment (Graham *et al*, 2019). Other antibodies may bind to non-essential molecules of the pathogen and provide limited protection (Caddy *et al*, 2021; Graham *et al*, 2019). For some infections, such as Severe Acute Respiratory Syndrome (SARS) virus and dengue virus, non-neutralizing antibodies may enhance infection and disease severity. In this process termed antibody-dependent enhancement (ADE), antibody-Fc receptor interactions promote uptake of the pathogen into Fc receptor-expressing immune cells (Graham *et al*, 2019; Wells *et al*, 2025). Hence, it is a priority to design molecular architectures that tailor immune responses to elicit the most protective antibodies, while minimizing the amount of deleterious antibody responses.

The earliest vaccines typically consisted of inactivated or attenuated pathogens, eliciting antibody responses to a range of pathogen-derived antigens (Morens *et al*, 2023). Subsequent advances enabled the development of more defined immunogens, including subunit vaccines, although this approach sometimes led to weaker and less sustained immune responses (Graham *et al*, 2019; Morens *et al*, 2023). Multivalent display of antigen subunits on virus-like particles (VLPs) provides a general strategy to combine the specificity of defined immunogens with enhanced immunogenicity. The larger size of VLPs promotes accumulation in lymph nodes, while the repetitive antigen arrangement facilitates B cell receptor crosslinking and enhances B cell activation (Bachmann & Jennings, 2010; Hills & Howarth, 2022). Structural biology of antibody-antigen recognition, either with monoclonal or polyclonal antibody samples, has provided great insight into features recognized by the immune system and how recognition correlates with efficacy, specificity and susceptibility to pathogen evasion (Graham *et al*, 2019; Sewall *et al*, 2025). Once a key protein antigen is identified, structure-based and computational approaches may be employed to improve its efficacy as a vaccine immunogen (Musunuri *et al*, 2024). Many methods have been tested to achieve immune focusing and orchestrate an antibody response that targets conserved antigens, to provide long-lasting protection to mutating viruses (Musunuri *et al*, 2024; Ward & Wilson, 2020). The antigen may be shortened, removing unwanted domains or truncating flexible regions (Graham *et al*, 2019). To ensure antibodies are raised to antigen in a relevant conformation, antigen stability may be enhanced with disulfide bonds, repacking the hydrophobic core, incorporation of rigidifying proline residues, or foldons to maintain the native antigen oligomerization state. In some cases, engineering can stabilize the antigen in the experimentally observed or putative pre-fusion state (Graham *et al*, 2019). N-linked glycans may be added or removed, to modify accessibility of the antigen surface to antibody induction (Joe *et al*, 2020; Zhang *et al*, 2022). Chemical coupling with low immunogenicity polymers like polyethylene glycol (PEGylation) or genetic fusion of repeating flexible peptide motifs (PASylation) has also been employed to conceal regions of antigens (Kraft *et al*, 2022; Weidenbacher & Kim, 2019). The orientation of antigens may be modified to enable the most important sites to project away from the surface of the vaccine platform (Frey *et al*, 2023). Different related antigens may also be given, either sequentially or simultaneously, such as through mosaic VLPs or tandemly linked repeats, directing the immune system to generate cross-reactive immune responses that recognize different strains of the same pathogen or even different related pathogens (Cohen *et al*, 2021; Hills *et al*, 2024; Ueda *et al*, 2020; Van Reeth *et al*, 2023).

Despite these advances, there is still a need for improved routes for molecular assembly of vaccines against major health challenges, including HIV, tuberculosis and dengue fever (Boomgarden & Upadhyay, 2025; Wang *et al*, 2024; Lee *et al*, 2025). There are important limitations in the level and specificity of antibodies that current vaccines elicit. Masking approaches may struggle to mask large surfaces, since each N-linked glycan only masks a small region of the antigen and high level glycosylation sometimes alters antigen folding or impairs antigen expression (Hyakumura *et al*, 2015; Ingale *et al*, 2014). Moreover, none of the immunofocusing methods described above actively reduce pre-existing unhelpful antibodies from past exposure. Here we take a distinct approach to guiding antibody induction, employing modular coupling of proteins and nanocage architectures, to generate a novel platform to directly decrease some immune responses while augmenting other responses. Inhibitory receptors are well known on T cells, leading to the breakthrough of checkpoint inhibitors in cancer therapy (Adashek *et al*, 2025). Inhibitory receptor signaling is also crucial to B cells, to avoid autoimmune responses (Tsubata, 2019). B cell activation involves increased activation of kinases relative to phosphatases adjacent to the B cell receptor (BCR) upon encountering a target antigen (Degn & Tolar, 2025). CD22 (also termed Siglec-2) is a B cell marker whose N-terminal domain recognizes sialic acid glycoconjugates and whose cytosolic C-terminus contains immunoreceptor tyrosine-based inhibition motifs (ITIMs) that engages tyrosine phosphatase signaling (Tsubata, 2019). If a B cell receptor (BCR) interacts with a self-antigen on an apposing cell-surface, CD22 recognizes terminal sialic acids across the immunological synapse and so recruited phosphatases turn down the stimulatory phosphorylation that would result from BCR cross-linking (Tsubata, 2019). FcγRIIb (also known as CD32B) is an inhibitory receptor that binds to the Fc region of IgG antibodies and recruits phosphatases through its ITIM, thus decreasing BCR signaling when there are already many antibodies recognizing the cognate antigen (Tsubata, 2019; Frampton *et al*, 2024). Stimulation of CD22 has been harnessed against autoimmunity using liposomes coated with sialic acid analogs, but there was no attempt to stimulate other immune responses at the same time (Macauley *et al*, 2013; Brzezicka *et al*, 2022).

Here we introduce Duality vaccination, which involves both a stimulatory immunogen and an inhibitory immunogen. The stimulatory immunogen induces responses to a particular pathogenic antigen, while the inhibitory immunogen drives decreased responses to antigenic sites where antibodies would not be helpful (Fig. 1A). We establish a framework to optimize this strategy with model antigens, bridging antigens with different antibodies to CD22 or FcγRIIb (Fig. 1B). Use of SpyTag003/SpyCatcher003 for high yield and modular amide bond formation allowed efficient construction of different immunogens (Fig. 1C) (Howarth, 2025), with assembly on the computationally-designed 60-mer mi3 nanocage for driving strong antibody induction (Bruun *et al*, 2018; Rahikainen *et al*, 2021). We demonstrate how we can immunofocus antibody responses as a function of valency, spacer organization and immunization timing, with potential wide impact for molecular engineering of potent immune responses.

**Fig. 1:**
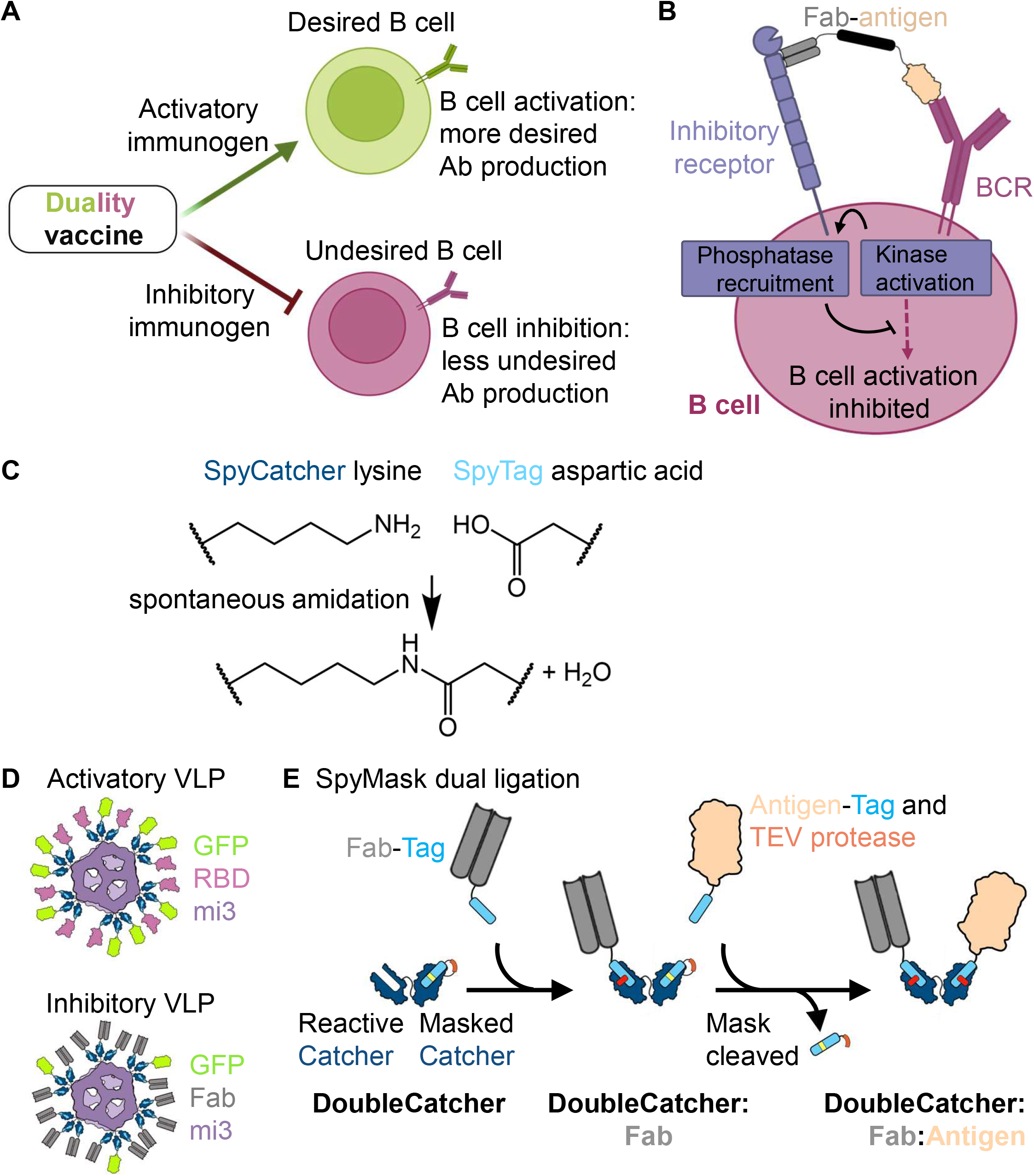
Introduction to Duality vaccination and immunogen nanoassembly. (**A**) Concept of Duality vaccination. A duality vaccine has an activatory immunogen which stimulates desired antibodies and an inhibitory immunogen to decrease the production of undesired antibodies. (**B**) Schematic of potential inhibitory mechanism. Linkage of the antigen with a binder to an inhibitory receptor (here shown with CD22) should lead to apposition of tyrosine phosphatases like SHP-1 near kinases like Lyn on the cytosolic tail of the B cell receptor (BCR), for inhibiting B cell activation. (**C**) Reaction between SpyTag and SpyCatcher moieties. (**D**) Illustration of activatory and inhibitory VLPs, with SpyTag003-linked antigen coupled to SpyCatcher003-mi3 nanocage. (**E**) Schematic of SpyMask assembly.

## Results

### Nanocage functionalization for duality immunization

We employed the Plug-and-Display approach for VLP functionalization, based upon a spontaneous amidation reaction between SpyCatcher003 on the mi3 VLP and SpyTag003 on the target antigens to be displayed (Brune & Howarth, 2018; Keeble & Howarth, 2020). The mi3 VLP is a computationally-designed 60-mer dodecahedral nanocage (Hsia *et al*, 2016), which we had previously modified to display SpyCatcher003 and optimized for stability and high yield expression from *E. coli* (Bruun *et al*, 2018; Rahikainen *et al*, 2021). Using SpyTag003/SpyCatcher003 coupling, it is straightforward to decorate these VLPs simultaneously with multiple antigens (Cohen *et al*, 2021; Hills & Howarth, 2022), serving as a model for a pathogen naturally displaying different antigens at its surface.

As a proof-of-principle, we employed SpyTag003-linked superfolder green fluorescent protein (GFP) and SARS-CoV-2 receptor-binding domain (RBD) as model antigens, which represent robust monomeric proteins allowing clear discrimination of antibody selectivity. We also recombinantly expressed Fab antibody fragments targeting two different B cell inhibitory receptors, each genetically fused with a SpyTag003 at the C-terminus of the heavy chain. Initial experiments were performed with Cy34.1 Fab which binds to mouse CD22 (Yu *et al*, 2015) and 128 or 130-2 Fabs which bind to mouse FcγRIIb with substantially different affinity (Williams *et al*, 2012). Fabs were expressed in Expi293F mammalian cells and purified via SpyTag003 using SpySwitch affinity chromatography (Vester *et al*, 2022). The purified Fabs were initially validated by SDS-PAGE with Coomassie staining and electrospray ionization mass spectrometry (Fig. S1). For further validation, we expressed soluble ectodomains of the cognate receptors in mammalian cells (Fig. S2). We then tested binding of the Fabs to their receptor by enzyme-linked immunosorbent assay (ELISA), quantifying affinity and binding kinetics by biolayer interferometry (BLI) (Fig. S3). These data confirmed that the 128 Fab had ∼5-fold lower affinity to FcγRIIb than the 130-2 Fab, as expected based on their parent monoclonal antibodies (Williams *et al*, 2012).

We constructed an activatory immunogen by assembling a VLP co-displaying GFP and RBD in equal molar amounts. In parallel, we created an inhibitory immunogen to turn down GFP-specific B cells, comprising a VLP co-displaying GFP and a Fab targeting an inhibitory receptor, in a 9:1 ratio (Fig. 1D). This design was analogous to previously published highly-multivalent Siglec-engaging tolerance-inducing antigenic liposomes (STALs) (Macauley *et al*, 2013). We confirmed uniform assembly of both unconjugated and conjugated VLPs by dynamic light scattering (DLS) (Fig. 2A). Precise covalent decoration of antigens and Fabs onto SpyCatcher003-mi3 was further determined by SDS-PAGE gel-shift analysis, following boiling in SDS to disrupt any non-covalent interactions (Fig. 2B).

**Fig. 2:**
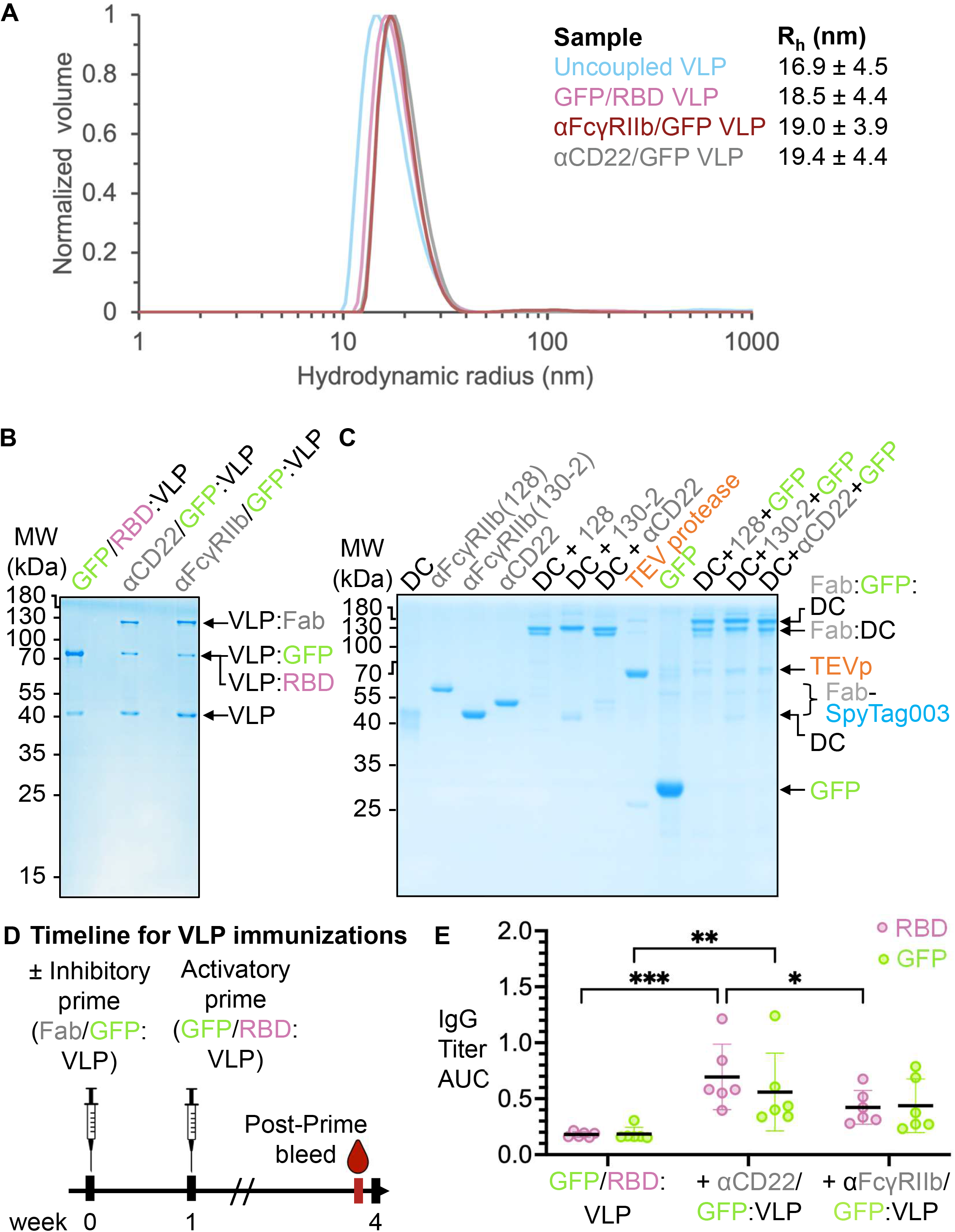
Immunizations using VLPs as both activatory and inhibitory components. (**A**) DLS for SpyCatcher003-mi3 (Uncoupled VLP), GFP/RBD VLP, anti-CD22/GFP VLP or anti-FcγRIIb/GFP VLP with hydrodynamic radius shown alongside. Representative curve shown (mean ± 1 s.d., n = 3). (**B**) Plug-and-Display assembly. SpyCatcher003-mi3 was mixed with SpyTag003-linked to GFP, RBD or Fab, before boiling in SDS and analysis by non-reducing SDS-PAGE with Coomassie staining. Molecular weight (MW) markers are indicated on the left. (**C**) Bispecific assembly with antigen and Fabs. Non-reducing SDS-PAGE with Coomassie staining to show DoubleCatcher (DC), MBP-TEV protease, and Fab building blocks, along with coupling to Fab alone, or coupling to Fab + antigen. (**D**) Immunization timeline, indicating the intramuscular injection of GFP/RBD VLPs with or without Fab/GFP VLPs, all without adjuvant, and serum sampling. (**E**) Antibody responses were enhanced by inhibitory receptor binders on the VLPs. ELISA for post-boost serum IgG against RBD (pink) or GFP (green) in response to activatory VLPs alone or with inhibitory VLPs containing Fabs against CD22 or FcγRIIb. Prime response from each mouse (n=6) is shown, expressed as IgG titer Area Under the Curve (AUC), with the black line showing the mean and error bars showing standard deviation. Statistical significance calculated using two-way ANOVA with Fisher’s Least Significant Difference (LSD) test. *p < 0.05, **p < 0.01, ***p < 0.001, no stars = non-significant.

Given the importance of receptor clustering in immune activation (Bachmann & Jennings, 2010; Davis & van der Merwe, 2006; Degn & Tolar, 2025), we hypothesized that the valency of our inhibitory immunogen would be a key determinant of its effect on immune activation. Therefore, in addition to the multivalent VLP inhibitory components, we generated using the SpyMask approach a bispecific construct containing a single antigen coupled to a single binder targeting an inhibitory receptor (Fig. 1E). SpyMask utilizes a DoubleCatcher (DC) construct containing two SpyCatcher003 moieties, one of which is available for reaction, while the second is masked by a non-reactive peptide (Driscoll *et al*, 2024). The first ligand bearing SpyTag003 may be coupled to DoubleCatcher, after which tobacco etch virus (TEV) protease releases the non-reactive SpyTag003 peptide and exposes the second SpyCatcher003 moiety for conjugation to a second ligand (Fig. 1E) (Driscoll *et al*, 2024). We used this modular approach to couple GFP-SpyTag003 at one site and a Fab against CD22 or FcγRIIb at the second site, generating bispecific constructs with 1:1 antigen to inhibitory receptor binder stoichiometry. We validated bispecific assembly by SDS-PAGE (Fig. 2C, Fig. S4A) and confirmed formation of both isopeptide bonds by electrospray ionization mass spectrometry (Fig. S4B-D).

### Inhibitory complexes on VLPs do not promote immunofocusing

We initially tested administration of the inhibitory VLPs one week prior to administering the activatory VLPs, to allow time for the inhibitory component to act before a stimulatory immune environment was generated. Published work reported that giving a CD22-inhibitory signal for 1 to 4 weeks prior to a stimulatory signal results in optimal inhibition (Macauley et al., 2013). Accordingly, we first injected mice intramuscularly with the VLP coupled to both GFP and an inhibitory Fab. One week later we injected the activatory VLP displaying both RBD and GFP (GFP/RBD VLP), hypothesizing that the resulting antibody response would be focused to the RBD immunogen. All injections were performed at the same site, without adjuvant. Four weeks after the initial injection we collected sera (Fig. 2D) and assessed the antigen-specific IgG antibody responses of the mice. We measured the IgG response to GFP or RBD by ELISA. Mice immunized only with GFP/RBD VLP showed minimal antibody responses to either RBD or GFP (Fig. 2E). In contrast, the response to both RBD and GFP was significantly higher in the mice that received the GFP/anti-CD22 VLP. The response to both RBD and GFP was also increased in mice that had received the GFP/anti-FcγRIIb VLP although this change did not reach statistical significance (Fig. 2E). Overall, VLPs without addition of adjuvant elicited minimal immune response, while co-displaying on the VLPs Fabs that target inhibitory BCRs non-specifically increased the overall immune responses. These results show that multivalency is so potent in enhancing immunogenicity (Bachmann & Jennings, 2010; Hills & Howarth, 2022) that multivalent presentation of GFP overcame ligation of inhibitory receptors and non-specifically enhanced the immune response.

### Testing bispecific immunogens with temporal and spatial separation from the activatory immunogen

We therefore focused on an inhibitory immunogen without high-level multimerization, assembled using DoubleCatcher (Fig. 1E). For immunizations with the DoubleCatcher bispecific inhibitory constructs, we tested whether the timing and site of administration of the Duality vaccine components influenced the response, since costimulatory signals generated within a highly inflammatory environment might override inhibitory signaling in B cells (Matsubara *et al*, 2018) (Fig. 3A & B). We also examined the effect of simultaneous delivery of activatory and inhibitory immunogens (prime responses in Fig. S5A and boost responses in Fig. 3C). If effective, simultaneous delivery would be advantageous as it would minimize the number of injections required.

**Fig. 3:**
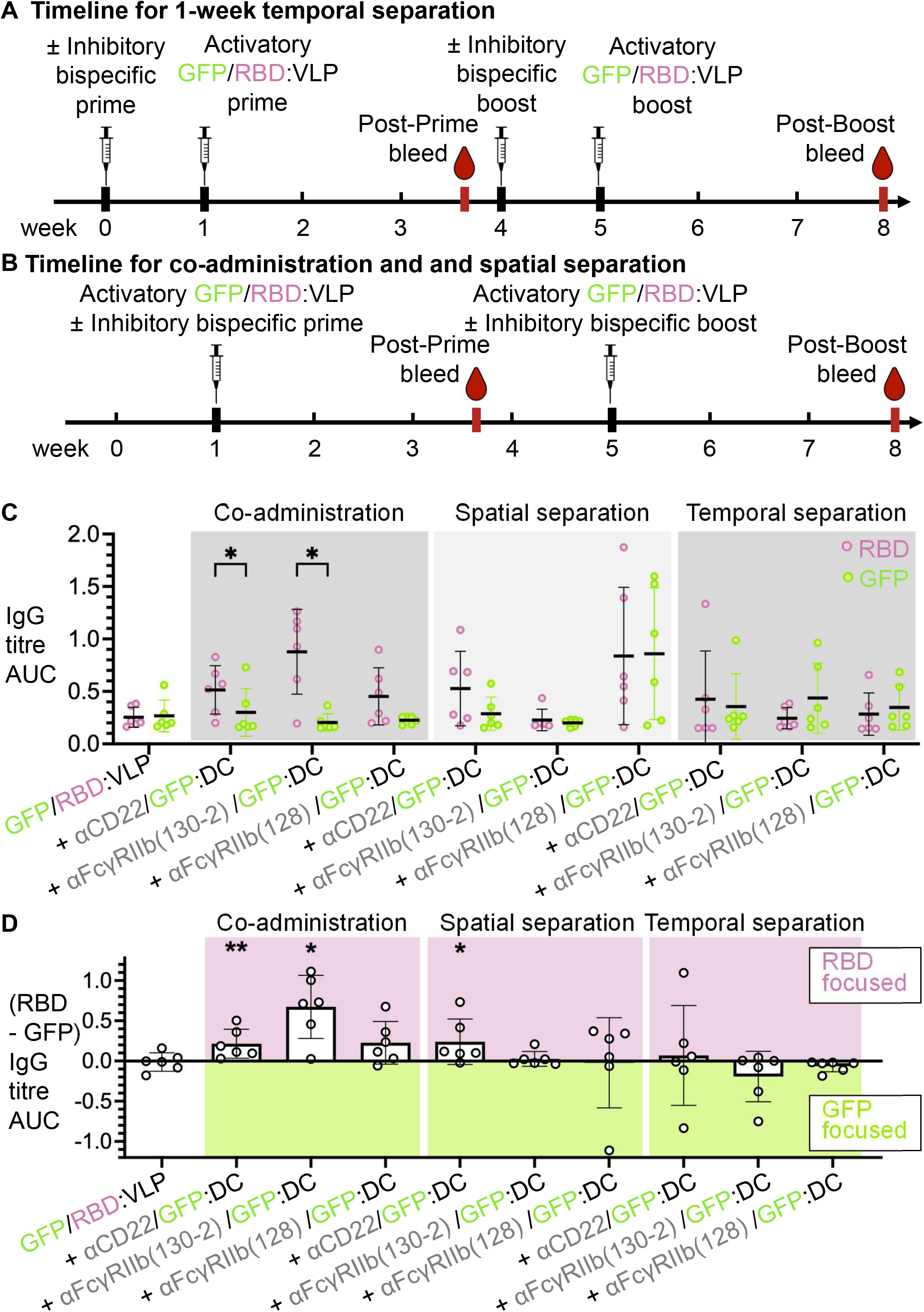
Temporal- and spatial-separation of activatory VLP and bispecific inhibitory immunogens. (**A**) Immunization timeline, indicating intramuscular injection of GFP/RBD VLPs ± Fab/GFP DoubleCatcher, all without adjuvant. With temporal separation, the inhibitory immunogen was first injected and then the activatory immunogen was injected one week later to the same leg. (**B**) Immunization timeline for co-administration or spatial separation (activatory and inhibitory immunogens injected on the same day to different legs). (**C**) ELISA for post-boost serum IgG against RBD (pink) or GFP (green) in response to activatory VLPs ± inhibitory DoubleCatcher containing Fabs against CD22 or FcγRIIb (130-2 or 128). Co-administration is shown in the left block, spatial separation in the center block, and temporal separation in the right block. Boost response from each mouse (n=6) is shown, with the black line showing the mean and error bars showing standard deviation. Statistical significance was calculated by paired t-tests. *p < 0.05, no stars = non-significant. (**D**) Summary of immunofocusing, based on (C). Titer against GFP was subtracted from titer against RBD. Dots represent single mice (n=6) with the block indicating the mean and error bars showing standard deviation. Statistical significance calculated by a t-test comparing each test group to GFP/RBD:VLP control. *p < 0.05, **p < 0.01, no stars = non-significant.

In all groups, mice were immunized intramuscularly with the activatory immunogen (GFP/RBD VLP) without adjuvant. In the co-administration groups, an inhibitory immunogen comprising GFP coupled to an inhibitory Fab through DoubleCatcher was mixed with the activatory immunogen before injection. For spatial separation, the activatory and inhibitory immunogens were injected into opposite gastrocnemius muscles (left and right leg) on the same day. For temporal separation, the inhibitory immunogen was administered one week before the activatory immunogen at the same site (Fig. 3A & B). The same regimens were used for boosting four weeks later. The IgG response to both RBD and GFP was weak following prime-boost immunization with the activatory immunogen alone. In contrast, co-administration of the activatory and inhibitory immunogens efficiently focused the IgG response towards RBD by generating a substantial anti-RBD response, while maintaining a weak anti-GFP response (Fig. 3C). The effect was most apparent with the high-affinity 130-2 Fab targeting FcγRIIb, although a significant immunofocusing was also observed with the anti-CD22 Fab (Fig. 3C, left panel). The difference between RBD and GFP response in groups receiving the lower-affinity 128 Fab targeting FcγRIIb did not reach significance (Fig. 3C, left panel). Plotting the difference between anti-RBD and anti-GFP titers for each mouse further highlighted the bias towards RBD in groups displaying successful immunofocusing (Fig. 3D). These results provided our first proof-of-concept that bridging an antigen to an inhibitory receptor can selectively decrease antigen-specific antibody responses during immunization.

The immunofocusing was not enhanced by either spatial separation or temporal separation compared with co-administration (Fig. 3C, middle and right panels). In fact, responses to both RBD and GFP appeared to be enhanced in the αFcγRIIb(128)/GFP:DC group following spatial separation. Given the lack of benefit from separating the vaccine components in either space or time, we therefore conducted subsequent Duality vaccination experiments via co-administration.

### Effects of adjuvant and immunogen dose on Duality vaccination

Although in some cases strong immune responses can be elicited by unadjuvanted SpyCatcher-linked VLPs (Brune *et al*, 2016), most vaccine regimens require adjuvant. Adjuvants provide strong co-stimulatory signals and recruit immune cells to the immunization site (Ben-Akiva *et al*, 2025). Adjuvants promote antigen uptake and presentation, enhancing germinal centre development and overall increasing the level and longevity of antibody responses. Adjuvant will also be essential to promote responses to low-immunogenicity antigens, such as in HIV (Ben-Akiva *et al*, 2025). To ensure the broad usefulness of our technology, therefore, we evaluated Duality vaccine immunofocusing in the presence of adjuvant.

As in the experiments without adjuvant, the activatory immunogen comprised GFP and RBD displayed on mi3 VLPs, while the inhibitory immunogen comprised GFP bridged through DoubleCatcher to Fabs targeting different inhibitory receptors. AddaVax was included as an adjuvant in a prime-boost regimen (Fig. 4A). In addition, we compared different doses of inhibitory immunogen, given the substantially greater immune response of multivalent VLPs relative to smaller immunogens (Bachmann & Jennings, 2010).

**Fig. 4:**
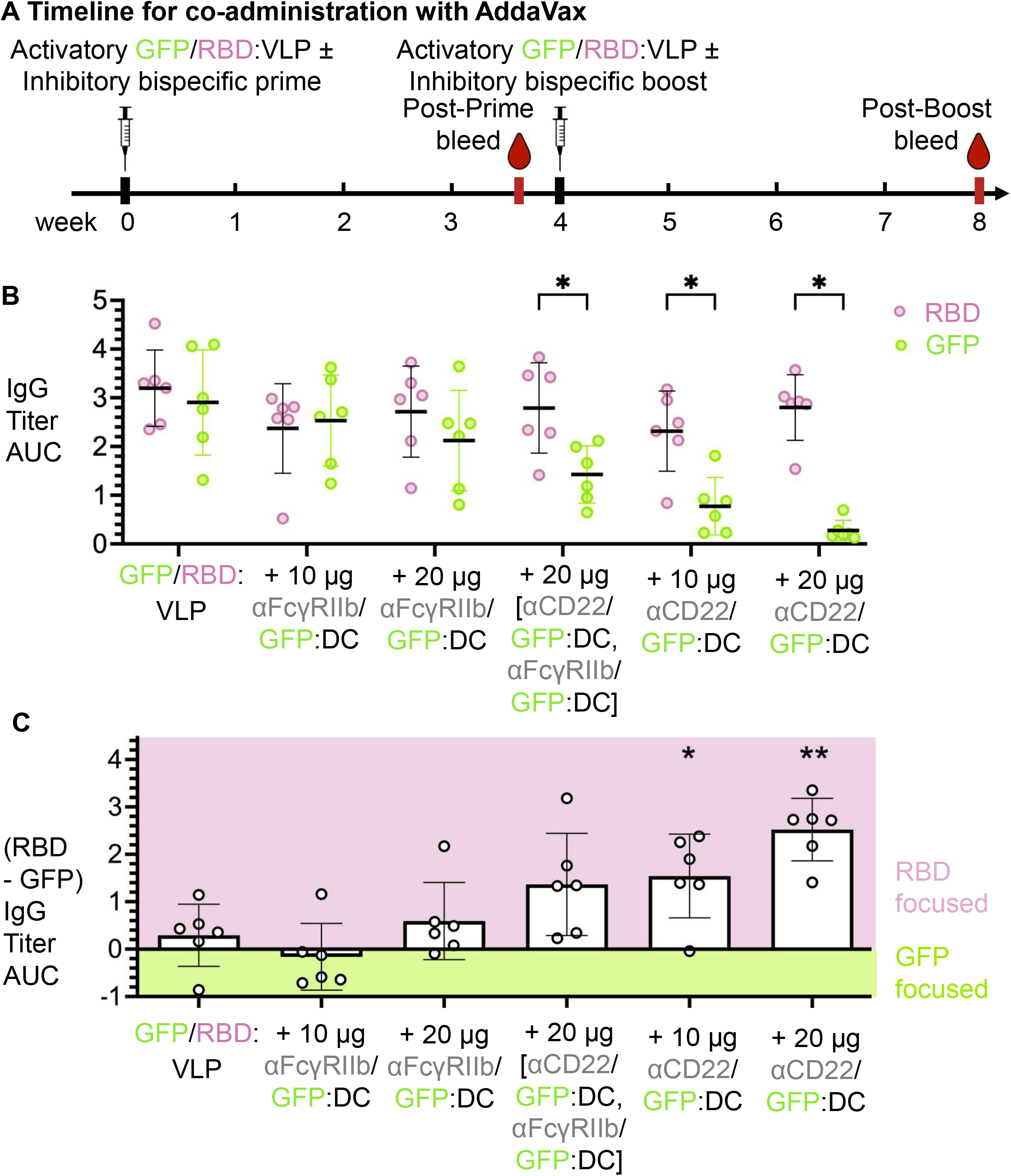
Duality vaccination with Adjuvant. (**A**) Immunization timeline, indicating intramuscular injection of Fab/GFP DoubleCatcher, co-administered with GFP/RBD VLPs and AddaVax adjuvant. (**B**) With adjuvant, anti-CD22 gave more immunofocusing than anti-FcγRIIb. ELISA for post-boost serum IgG against RBD (pink) or GFP (green) in response to activatory VLPs ± inhibitory DoubleCatcher containing Fabs against CD22 or FcγRIIb (130-2). Dose refers to µg of GFP on the inhibitory component. For the group receiving both αCD22/GFP:DC and αFcγRIIb(130-2)/GFP:DC, the 20 µg dose is made up of 10 µg each. Boost response from each mouse (n=6) is shown, with the black line showing the mean and error bars showing standard deviation. Statistical significance was calculated by paired t-tests. *p < 0.05, no stars = non-significant. (**C**) Summary of immunofocusing, based upon (B). Titer against GFP was subtracted from titer against RBD, to highlight the different focusing. Dots represent single mice (n=6) with the block indicating the mean and error bars showing standard deviation. Statistical significance calculated by a t-test comparing each test group to GFP/RBD:VLP control. *p < 0.05, **p < 0.01, no stars = non-significant.

Post-boost antibody titers were an order of magnitude higher than in previous experiments without adjuvant. Strikingly, in the presence of adjuvant, immunofocusing away from GFP was more pronounced in experiments with the anti-CD22 Fab than with the anti-FcγRIIb 130-2 Fab (Fig. 4B & C). Targeting CD22 did not significantly alter the IgG response to RBD but greatly reduced the response against GFP (Fig. 4B & C). The decrease of the anti-GFP response was stronger with a higher dose of inhibitory anti-CD22 immunogen (Fig. 4B & C; prime responses in Fig. S5B). We also investigated whether combining anti-CD22 and anti-FcγRIIb inhibitory immunogens could enhance immunofocusing, but the combination provided no additional improvement (Fig. 4B & C). Together, these results demonstrate that Duality vaccination can achieve selective immunofocusing in the presence of adjuvant, and that, under these conditions, monovalent targeting of CD22 was more effective than targeting FcγRIIb or combining the two inhibitory immunogens.

### Linker-dependence of Duality vaccination

Having successfully and selectively decreased the antibody response to GFP, we next assessed whether Duality vaccination could similarly allow selective decrease in the response to RBD. Having established the optimal inhibitory activity of the anti-CD22 Fab in the preceding experiments, we assembled this Fab with RBD using DoubleCatcher and validated the assembled construct by SDS-PAGE (Fig. 5A). We then performed immunizations with the activatory immunogen (GFP/RBD VLP) together with inhibitory immunogens comprising anti-CD22 Fab linked to either GFP or RBD, following the dosing schedule in Fig. 4A. Co-administration of anti-CD22/GFP again strongly reduced the anti-GFP response (boost responses in Fig. 5B; prime responses in Fig. S5C). More direct bridging, using anti-CD22 Fab directly fused to SpyCatcher003 and coupled to GFP-SpyTag003, also successfully decreased the anti-GFP antibody response. However, anti-CD22/RBD had no detectable effect on the antibody response to RBD (Fig. 5B & H).

**Fig. 5:**
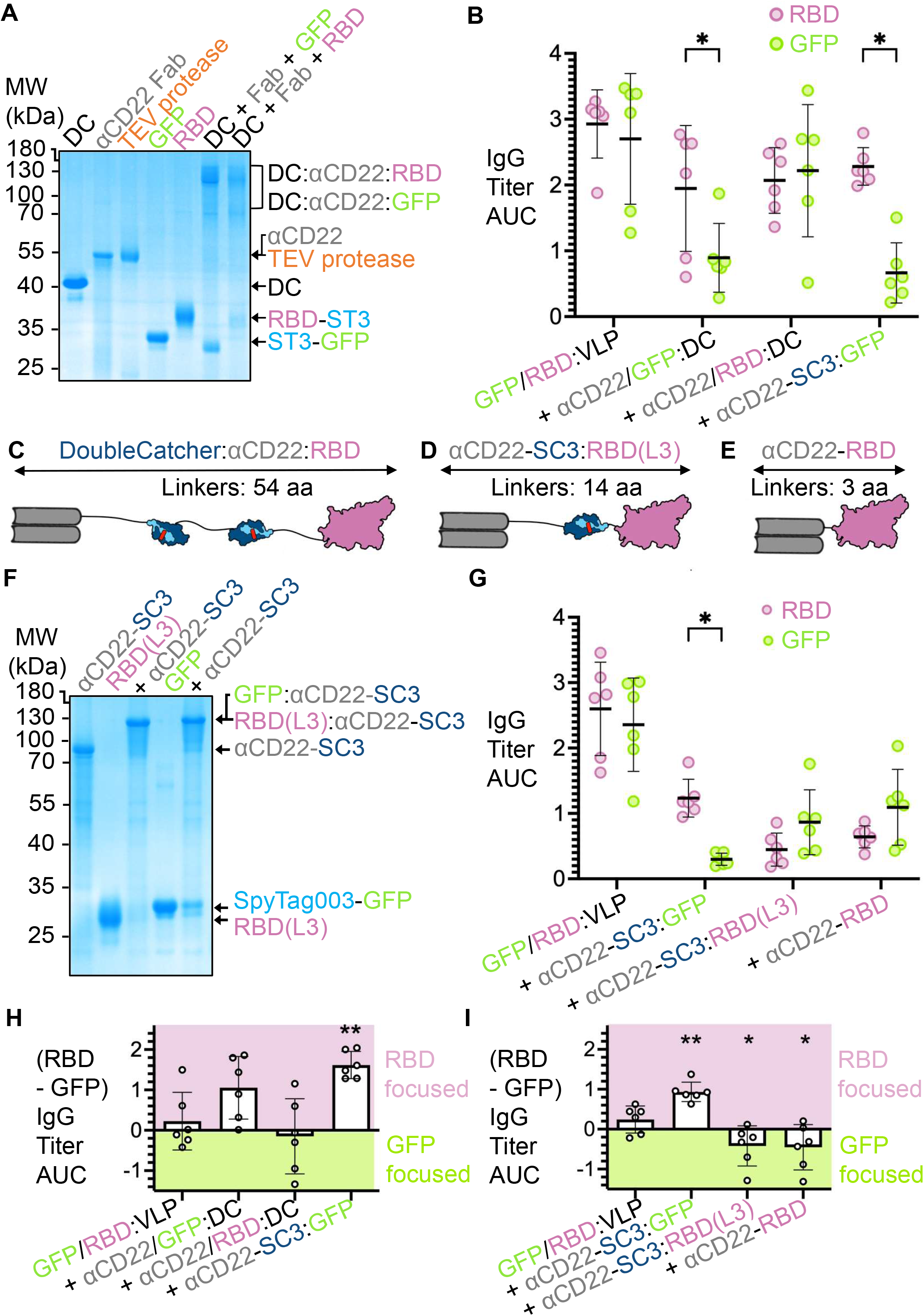
Linker dependence for Duality vaccination. (**A**) Assembly of inhibitory immunogen using SpyMask, assessed by non-reducing SDS-PAGE with Coomassie staining. (**B**) Biased response from inhibitory immunogen based on GFP but not based on RBD. ELISA for post-boost serum IgG against RBD (pink) or GFP (green) in response to prime and boost with adjuvant using activatory VLPs alone or with inhibitory DoubleCatcher containing anti-CD22 Fab and GFP or RBD or anti-CD22-SpyCatcher003:GFP. Boost response from each mouse (n=6) is shown, with the black line showing the mean and error bars showing standard deviation. Statistical significance was calculated by paired t-tests. *p < 0.05, no stars = non-significant. (**C, D, E**) Schematic of inhibitory constructs. Structural representations, based on AlphaFold3 and PyMOL visualization, illustrating the long linker on DoubleCatcher:anti-CD22:RBD (C) compared to anti-CD22-SpyCatcher003:RBD(L3) (D) or the direct genetic fusion anti-CD22-RBD (E). Total number of amino acid residues (aa) in flexible Glycine-Serine linkers is given for each construct. (**F**) Assembly of inhibitory immunogen by direct coupling with anti-CD22 Fab genetically fused to SpyCatcher003, assessed by non-reducing SDS-PAGE with Coomassie staining. **(G**) Change in antibody bias according to spacer length. ELISA for post-boost serum IgG against RBD (pink) or GFP (green) in response to prime and boost with adjuvant using activatory VLPs alone or with inhibitory immunogens with anti-CD22-SpyCatcher003 linked to GFP or RBD(L3) or anti-CD22-RBD fusion. Boost response from each mouse (n=6) is shown, with the black line showing the mean and error bars showing standard deviation. Statistical significance was calculated by paired t-tests. (**H**) and (**I**) Summary of immunofocusing, based upon (B) and (G), respectively. The antibody titer against GFP was subtracted from the titer against RBD, to highlight the different focusing. Dots represent single mice (n=6) with the block indicating the mean and error bars showing standard deviation. Statistical significance calculated by a t-test comparing each test group to GFP/RBD:VLP control. *p < 0.05, **p < 0.01, no stars = non-significant.

We considered several possible explanations for the difference in effects of the GFP- or RBD-targeted inhibitory immunogens. One notable difference was the spacing between antigen and SpyTag003, since the RBD construct contained a long 34-residue linker to SpyTag003, whereas the GFP construct contained an 11-residue linker. Our initial constructs were designed with glycine/serine-rich spacers, imparting flexibility and promoting efficient coupling to SpyCatcher003 (Keeble & Howarth, 2019). However, proximity between the phosphatases associated with inhibitory receptors and the BCR is likely to be important for turning down the overall B cell response (Tsubata, 2019). We hypothesized that the greater separation allowed by the RBD construct might limit its ability to inhibit B cell activation. To investigate the effect of spacing, we generated a series of inhibitory immunogens with progressively reduced linker lengths between the anti-CD22 and RBD. First, we expressed a new RBD construct in which the linker to SpyTag003 contained only three amino acid residues, named RBD(L3). We also further reduced the number of linking residues using a simplified inhibitory immunogen architecture, in which SpyCatcher003 was directly fused to the C-terminus of the Fab heavy chain for coupling to SpyTag003-linked antigen (Fig. 5D & F). Finally, we generated a direct fusion of anti-CD22 Fab to RBD, taking advantage of the efficient mammalian expression of both components (Fig. 5E and S6A). The architectures of these constructs, with their linkers depicted in extended conformation, are illustrated in Fig. 5C-E and Fig. S6B.

Immunization with the shorter inhibitory immunogens resulted in selective inhibition of the response to RBD, such that despite some reduction in the antibody GFP response, the response to GFP was now higher than the response to RBD (Fig. 5G; prime responses are shown in Fig. S5D). Plotting the difference between anti-RBD and anti-GFP titers highlighted that the direction of immunofocusing could be shifted towards either RBD or GFP, according to the inhibitory immunogen selected (Fig. 5H & I). Thus, Duality vaccination could selectively decrease antibody responses to either a bacterially-expressed, non-glycosylated antigen (i.e. GFP) or a mammalian-expressed virus-derived glycosylated antigen (i.e. RBD). These findings further demonstrated that the structural organization, particularly the linker design of the inhibitory immunogen, is a crucial determinant of effective immunofocusing.

### Immunofocusing within a large antigen

Having demonstrated that Duality vaccination could selectively decrease responses to individual antigens, we next assessed whether this approach could reshape immunodominance between different epitopes of a larger antigen. Spike protein is the dominant target of natural neutralizing antibody responses following SARS-CoV-2 infection (Kleanthous *et al*, 2021). Spike’s extracellular portion comprises the S1 and S2 subunits, with S1 containing the N-terminal domain (NTD) and RBD (Fig. 6A). The sequence conservation of these regions (Fig. 6B, Fig. S7) and their relative importance for virus neutralization have been extensively studied (Kleanthous *et al*, 2021), with some reports suggesting that focusing the response away from RBD and towards the more conserved S2 region may lead to more cross-protective responses (Guenthoer *et al*, 2024). To investigate the ability of the Duality vaccine strategy to alter the immunodominance hierarchy within Spike, we expressed and purified a stabilized HexaPro version of SARS-CoV-2 spike (Hsieh *et al*, 2020) in Expi293F cells (Fig. 6C). Soluble RBD, S2 and NTD subunits were also expressed separately in Expi293F cells (Fig. 6C). We used an inhibitory immunogen (anti-CD22:RBD) based upon anti-CD22-SpyCatcher003 covalently coupled to SARS-CoV-2 RBD(L3)-SpyTag003, bearing the short linker from antigen to SpyTag003 (as in Fig. 5D). All immunizations were performed with AddaVax adjuvant. Mice were primed and boosted with either Spike alone, or Spike together with anti-CD22:RBD(L3) (Fig. 6D). Antibody responses against full-length Spike, S2, NTD or RBD were measured by ELISA (boost responses in Fig. 6E; prime responses in Fig. S5E).

**Fig. 6:**
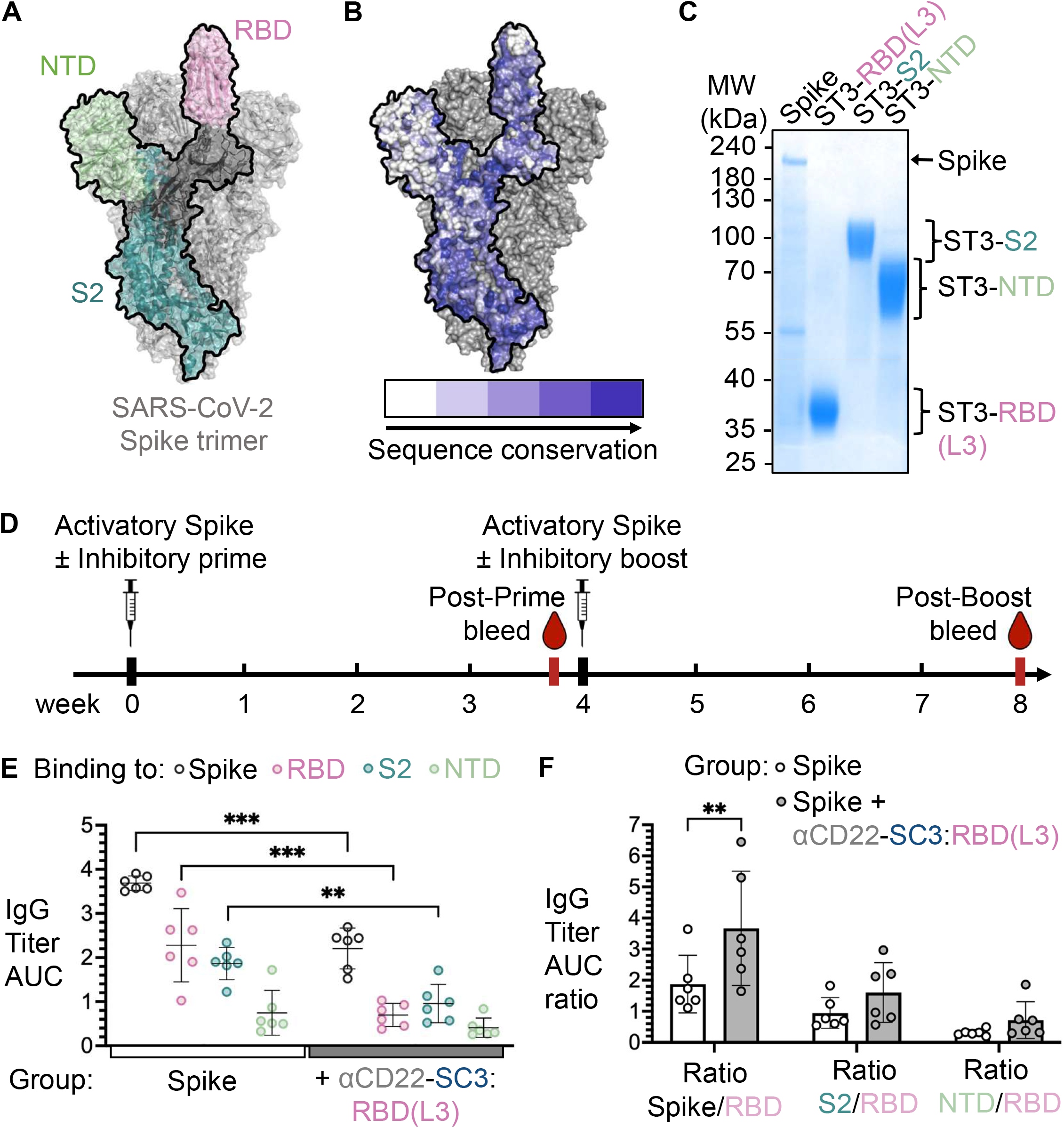
Immune focusing within a large viral antigen. (**A**) Regions of SARS-CoV-2 Spike ectodomain. One monomer of the trimer is outlined in black. The N-terminal domain (NTD) is shown in green, the S2 subunit in teal, and the receptor-binding domain (RBD) in pink, based on PDB 6VSB. (**B**) Sequence conservation in the Spike, based on 32 Coronavirus Spike proteins, color-coded from white as highly variable to dark blue highly conserved, mapped onto SARS-CoV-2 Spike ectodomain. (**C**) Purification of SpyTag003-linked SARS-CoV-2 Spike and RBD, S2 and NTD subunits, shown by SDS-PAGE with Coomassie staining. (**D**) Immunization timeline, indicating the prime and boost intramuscular injection of Spike and adjuvant, with or without anti-CD22-SpyCatcher003:RBD(L3) inhibitory immunogen. (**E**) ELISA for post-boost serum IgG against Spike (gray), RBD (pink), S2 (teal) or NTD (green) in response to activatory Spike alone (left) or with anti-CD22-SpyCatcher003:RBD(L3) (right). Boost response from each mouse (n=6) is shown, with the black line showing the mean and error bars showing standard deviation. Statistical significance for each domain was calculated using a two-way ANOVA with Fisher’s Least Significant Difference (LSD). (**F**) Summary of immunofocusing, based upon (E). Titer against Spike (left), S2 (middle) or NTD (right) was divided by titer against RBD, to highlight different focusing. Dots represent single mice (n=6) with the block indicating the mean and error bars showing standard deviation. Statistical significance calculated by a t-test. *p < 0.05, **p < 0.01, no stars = non-significant.

Following prime-boost immunization with Spike alone, the antibody response was strongest against the RBD domain compared to the other regions of Spike, whereas the response against NTD was weak (Fig. 6E), consistent with previous studies (Kleanthous *et al*, 2021). Co-administration of Spike with anti-CD22-SpyCatcher003:RBD(L3) shifted the immunodominance pattern. The relative response to RBD was reduced; S2 instead became the region eliciting the highest antibody response (Fig. 6E). To quantify this redistribution, we compared responses to each region relative to the response to full Spike for individual mice (Fig. 6F). There was a significant increase in the ratio of response to Spike compared to response to the RBD domain, but the increase in S2/RBD ratio and NTD/RBD ratio did not reach statistical significance (Fig. 6F). These data indicate that Duality vaccination can redirect immunodominance within a large, glycosylated and immunologically important viral antigen.

## Discussion

Here we establish Duality vaccination as a strategy for focusing antibody responses through the coordinated activation and inhibition of antigen-specific B cells. The approach combines multivalent antigen display on protein nanocages to enhance immune activation with click biology-based recruitment of inhibitory receptors to selectively decrease responses against defined antigenic features. Targeting CD22 was more effective than targeting FcγRIIb in the presence of adjuvant. Immune inhibition was highly sensitive to molecular architecture: efficient immunofocusing required the linker to be short between the antigen and the anti-CD22 targeting domain. High level multimerization of anti-CD22 Fab on a VLP abolished the inhibitory effect. These findings identify receptor choice, molecular spacing, and valency as key determinants of immunofocusing by Duality vaccination.

To characterize immunofocusing precisely, we employed SARS-CoV-2 RBD and GFP as model antigens. Both antigens represent two monomeric proteins suitable for evaluating antibody responses with high solubility, but differing in one being virus-derived and N-glycosylated (Watanabe *et al*, 2020), whereas the other has no post-translational modification other than the autocatalytic amino acid modifications that confer fluorescence (Tsien, 1998). Interestingly, we found that the immune response to GFP was more readily decreased than that to RBD. Future work may explore how antigen glycosylation, which may affect interaction with CD22 or other Siglecs (Macauley *et al*, 2013), or other features of the germline antibody repertoire may contribute to that difference. Notably, specific immune inhibitory effects were minimal following priming and became pronounced only after boosting.

Other inhibitory receptors such as PD-1 are also expressed on B cells (Thibult *et al*, 2013), but targeting PD-1 might have broader consequences in other lymphocyte populations. In particular, modulation of PD-1 could interfere with T cell activation that may be important for immunity against the activatory immunogen. B cells are sophisticated in their integration of multiple signals in the decision to mount an antibody response (Tsubata, 2019). In particular, pathogen-associated molecular patterns or pro-inflammatory signals, as enhanced by adjuvant, have important effects on the threshold and pattern for cell activation (Matsubara *et al*, 2018). Thus, it was an important finding that we could retain immunofocusing in the presence of the potent adjuvant AddaVax.

The different effects of CD22 and FcγRIIb targeting may reflect differences in their signaling architecture. The cytosolic tail of CD22 contains three ITIMs, as well as an ITIM-like motif and a Grb2-binding motif (Bornhöfft *et al*, 2018), whereas FcγRIIb contains a single ITIM (Roghanian *et al*, 2018). Future work may determine how this different modular organization may relate to the distinct response that we find upon co-ligation with the BCR of these different inhibitory receptors. Recruitment of CD22 to the BCR would not be expected to constitute an absolute block to B cell activation, since viral glycoproteins are often sialylated and protective antibody responses must still be mounted against such targets (Bornhöfft *et al*, 2018). Consistent with this principle, high-level multimerization of anti-CD22 Fab on VLPs failed to diminish the antibody response. We propose that the strong BCR clustering induced by multivalent antigen display is sufficient to overcome CD22 recruitment and allow strong BCR activation (Hills & Howarth, 2022). We hypothesize that avoiding extensive BCR clustering by the inhibitory immunogen may be important for Duality vaccination. We achieved this controlled architecture using the three-component SpyMask strategy or by a simpler two-component SpyTag003-antigen ligation to Fab-SpyCatcher003. This approach differs from previous strategies in which multivalent low affinity ligands displayed on liposomes recruited CD22 to induce antigen-specific tolerance in the absence of adjuvant (Macauley *et al*, 2013; Brzezicka *et al*, 2022). In contrast, here we: (i) used a nanomolar affinity Fab to recruit CD22 in close proximity to the BCR, and (ii) included adjuvant to help priming of new immune responses. Following our proof-of-concept study using SpyMask modular assembly to identify an effective bridging antibody, we showed that direct fusion of antigen to anti-CD22 was sufficient for immunofocusing, as long as the post-translational modifications of the two components were compatible. There are precedents for high sensitivity to spacer size for receptor signal transduction (Mohan *et al*, 2019; Fernandes *et al*, 2020). Hence there may be an optimal radius for Duality vaccine effect. However, this radius may differ between antigens of different dimensions and between BCRs recognizing different epitopes on specific antigens. Hence we have built modules of different size and retained flexibility in our linkers. If the BCR recognizes a region on a large antigen many nanometers from the anti-CD22 module, the inhibitory effect may be lessened.

Substantial challenges remain in directing vaccine-induced antibody responses, particularly for pathogens with a high degree of variability or immune evasion, such as HIV (Haynes *et al*, 2023). Duality vaccination provides a complementary approach to existing immunofocusing strategies and could potentially be combined with methods such as computationally-modified antigen scaffolds (Castro *et al*, 2025) or engineered glycan shielding (Musunuri *et al*, 2024). Future work may identify how Duality vaccination depends upon different routes of administration and translates to different species, towards urgent global priorities for both human and veterinary vaccines.

## Acknowledgements

G.A., R.A.H. and C.L.D. disclose support from the David James Trust Fund. M.R.H. acknowledges University of Cambridge start-up funds. R.A.H. discloses support for the research of this work from the Rhodes Trust. C.L.D. acknowledges funding by the Biotechnology and Biological Sciences Research Council (BBSRC) Oxford Interdisciplinary Bioscience Doctoral Training Partnership (DTP) (grant number BB/T008784/1). R.A.H. and T.K.T. disclose support for the research of this work from the Townsend-Jeantet Prize Charitable Trust (Charity Number 1011770). T.K.T. and D.M. disclose support for the research of this work from the Coalition for Epidemic Preparedness Innovations grant under the broadly protective coronavirus (BPCV) portfolio. We thank the Biophysics Facility and the Centre for Experimental Biophysical Chemistry at the University of Cambridge for assistance. P.J.B. discloses support for the research of this work from the Caltech Merkin Institute and a George Mason University Fast Grant. Research reported in this publication was supported by the National Institute of Allergy and Infectious Diseases of the National Institutes of Health under Award Number AI165075. The content is solely the responsibility of the authors and does not necessarily represent the official views of the National Institutes of Health. This manuscript is the result of funding in whole or in part by the National Institutes of Health (NIH). It is subject to the NIH Public Access Policy. Through acceptance of this federal funding, NIH has been given a right to make this manuscript publicly available in PubMed Central upon the Official Date of Publication, as defined by NIH. M.S.C. acknowledges support from Cancer Research UK (CRUK) programme DRCRPG-May23/100001. For the purpose of Open Access, the author has applied a CC BY public copyright licence to any Author Accepted Manuscript (AAM) version arising from this submission.

## Author Contributions Statement

G.A. and R.A.H. performed all experiments except C.L.D. generated DoubleCatcher assemblies and ran mass spectrometry, and T.K.T. and D.M. performed mouse immunizations. G.A., R.A.H., A.A.C., M.S.C., P.J.B., T.K.T. and M.R.H. designed the project. G.A. and M.R.H. wrote the manuscript. All authors read, reviewed and approved the manuscript.

## Methods

### Plasmids and Cloning

Constructs were cloned by PCR using Q5 High-Fidelity Polymerase (New England Biolabs). Fragments were assembled by Gibson assembly. Inserts of all constructs were validated by Sanger Sequencing (Source Bioscience).

The following plasmids have previously been published: pET28a-SpyCatcher003-mi3 (GenBank MT945417, Addgene 159995) (Rahikainen *et al*, 2021), pET28a-SpyTag003-MBP (GenBank MN433888, Addgene 133450) (Keeble *et al*, 2019), pET28a-SpyTag003-GFP (GenBank ON131080, Addgene 133454) (Keeble *et al*, 2019), pDEST14-SpyCatcher003-(GSG)_3_-SpyCatcher003-TEVs-SpyTag003DA (DoubleCatcher) (GenBank PP341218, Addgene 216284) (Driscoll *et al*, 2024), pET28-SpyCatcher002-MBP (GenBank PP341225, Addgene 216295) (Driscoll *et al*, 2024), pDEST14-SpyCatcher003 S49C (Addgene 133448) (Vester *et al*, 2022), pGEX-GST-BirA (glutathione-S-transferase-BirA) (O’Callaghan C *et al*, 1999) and pET28-MBP-superTEV protease (Addgene 171782) (Keeble *et al*, 2022). GST-TEV protease in pMinVecK was purchased from DNASU (ID TvCD00541083, a kind gift of Paul Blommel and Brian Fox). pOPH6 for Peptide:N-glycosidase F (PNGase F) expression was a kind gift from Shaun Lott (Addgene plasmid ID 40315).

RBD from SARS-CoV-2 Wuhan (residues 319-537) (Vester *et al*, 2022) (GenBank ON131086) contains a C-terminal His_8_-tag followed by a SpyTag003 (RGVPHIVMVDAYKRYK) (Keeble *et al*, 2019) to give p3BNC-RBD-His8-SpyTag003. The total length between the end of the RBD sequence and the start of SpyTag003 is 34 residues. RBD(L3) (pcDNA3.1-SpyTag003-GSG-RBD) was designed with only a 3 residue GSG linker between an N-terminal SpyTag003 and SARS2 RBD (residues 331-529), followed by a C-terminal His_6_ tag (GenBank and Addgene deposition in progress). The stabilized HexaPro SARS2 Wuhan Spike protein, a kind gift from Jason McLellan, was previously published (Hsieh *et al*, 2020) (Addgene 154754). pcDNA3.1-SpyTag003-S2 was constructed by cloning residues 686-1208 from Wuhan Spike into pcDNA3.1 with an N-terminal SpyTag003 (GenBank and Addgene deposition in progress). Similarly, pcDNA3.1-SpyTag003-NTD was constructed to encode Spike N-terminal domain (residues 13-303) (GenBank and Addgene deposition in progress).

Sequences for the anti-mouse CD22 Fab based on Cy34.1.2 monoclonal antibody were kindly provided by Henri-Jacques Delecluse (German Cancer Research Center, DKFZ) (Yu *et al*, 2015). Sequences were synthesized (IDT) and cloned into pcDNA3.1, with a C-terminal SpyTag003 for the heavy chain, giving pcDNA3.1-αCD22(Heavy)-SpyTag003 which has an 11 residue GS linker between the Fab and SpyTag003 (GenBank and Addgene deposition in progress) and pcDNA3.1-αCD22(Light) (GenBank and Addgene deposition in progress). pcDNA3.1-αCD22(Heavy)-SpyCatcher003-His_6_ was constructed by replacing SpyTag003 with SpyCatcher003 and adding a C-terminal His_6_, with an 11-residue GS linker between the Fab and SpyCatcher003 (GenBank and Addgene deposition in progress). pcDNA3.1-αCD22(Heavy)-RBD (GenBank and Addgene deposition in progress) has a 3-residue GSG linker between the Fab and RBD domains, which was constructed from pcDNA3.1-SpyTag003-GSG-RBD and pcDNA3.1-αCD22(Heavy)-SpyTag003-His.

Fab portions from anti-mouse FcγRIIb monoclonal antibodies AT-130-2 and AT-128 (Williams *et al*, 2012) were synthesized (IDT) and cloned into pcDNA3.1, giving pcDNA3.1-AT130-2(Heavy)-SpyTag003, which has a 13-residue GS linker between the Fab and SpyTag003 (GenBank deposition in progress). This cloning yielded pcDNA3.1-AT130-2(Light) (GenBank deposition in progress), pcDNA3.1-AT128(Heavy)-SpyTag003, which has a 15-residue GS linker between the Fab and SpyTag003 (GenBank deposition in progress), and pcDNA3.1-AT128(Light) (GenBank deposition in progress). To make pcDNA3.1-mCD22-His-AviTag (GenBank and Addgene deposition in progress), domains 1-3 of mouse CD22 (residues 1-347) were synthesized (IDT) and cloned into pcDNA3.1 with a C-terminal His_6_ tag and AviTag. pDSG102-mCD32-AviTag (GenBank and Addgene deposition in progress) contains murine FcγRIIb (mCD32 Ly17.2) extracellular portion (residues 29-207), based on pDSG102-mCD32_Ly17.2-His6.

### Bacterial expression

pET28a or pDEST14 plasmids were transformed into *E. coli* BL21(DE3) from Agilent by heat shock transformation. The cells were plated onto LB-agar plates containing either 50 μg/mL kanamycin (pET28a) or 100 µg/mL ampicillin (pDEST14) and grown overnight at 37 °C. A colony was picked to inoculate 20 mL LB with the appropriate antibiotic and grown for 16 h at 37 °C with shaking at 200 rpm. This starter culture was used to inoculate 1-4 L of LB with kanamycin or ampicillin and grown at 37 °C with shaking at 200 rpm until culture density reached OD_600_ 0.6-0.7. Cultures were induced by adding 0.5 mM isopropyl β-D-1-thiogalactopyranoside (IPTG). After 4 h at 30 °C with shaking at 200 rpm (for DoubleCatcher, SpyTag003-GFP and SpyTag003-MBP, SpyCatcher003 S49C, SpyCatcher002-MBP, MBP-superTEV and GST-TEV) or 16 h at 22 °C shaking at 200 rpm (for SpyCatcher003-mi3), cells were pelleted by 15 min centrifugation at 4,000 g.

pGEX-GST-BirA was expressed in *E. coli* BL21 (DE3) RIPL and purified using glutathione-sepharose as described (Fairhead & Howarth, 2015).

### Purification of SpyCatcher003-mi3

SpyCatcher003-mi3 was purified by ammonium sulfate precipitation (Rahikainen *et al*, 2021). Bacterial pellets were resuspended in 20 mM Tris-HCl, 300 mM NaCl, pH 8.5, supplemented with 0.1 mg/mL lysozyme, 1 mg/mL cOmplete mini EDTA-free protease inhibitor (Roche) and 1 mM phenylmethanesulfonyl fluoride (PMSF). This resuspended pellet was incubated for 30 min at 25 °C with end-over-end mixing. Sonication was then performed on ice with an Ultrasonic Processor equipped with a microtip (Cole-Parmer), for four rounds of 60 s on 50% duty-cycle, separated by a 60 s pause. This crude lysate was then centrifuged for 45 min at 35,000 *g* at 4 °C to remove cell debris. Then, 170 mg ammonium sulfate per mL of cleared lysate was added little by little, before incubating for 1 h with 120 rpm mixing from a magnetic stirrer. Precipitated particles were pelleted by centrifugation at 30,000 g for 30 min at 4 °C. The protein pellet was resuspended in 8 mL mi3 buffer (25 mM Tris-HCl, 150 mM NaCl, pH 8.0) at 4 °C and filtered through 0.45 µm and 0.22 µm syringe filters (Starlab). The sample was dialyzed for 16 h against 5,000-fold excess of the same buffer in a dialysis tube with 100 kDa cutoff, then centrifuged at 17,000 g for 30 min at 4 °C to pellet any insoluble material. Size-exclusion chromatography was then performed using a HiPrep Sephacryl S-400 HR 16-60 column (GE Healthcare) run on an ÄKTA Pure 25 system (GE Healthcare). The sample was run at 4 °C in mi3 buffer and collected in 1 mL elution fractions. Fractions containing purified particles as determined by A_280_ absorbance were pooled and concentrated using a Vivaspin 20 100 kDa molecular weight cut-off centrifugal concentrator (GE Healthcare) and stored at - 80 °C.

### Mammalian protein expression

p3BNC, pcDNA3.1 and pDSG102 constructs were expressed using Expi293F cells (Thermo Fisher). Expi293F cells were grown at 37 °C and 8% (v/v) CO_2_ in Expi293 Expression Medium (Thermo Fisher). Transfections were performed using the ExpiFectamine 293 Transfection Kit (Thermo Fisher), following manufacturer’s instructions. Briefly, 1 μg DNA per mL culture to be transfected was incubated with ExpiFectamine293 reagent for 10 min, then added to Expi293F cells at a density of 3 million cells per mL. After 18–22 h, ExpiFectamine 293 Transfection Enhancers 1 and 2 were added. After 5-7 days, supernatant was harvested by spinning cultures for 10 min at 4,000 g and filtered through a 0.22 μm filter (Starlab).

### Nickel affinity purification

His-tagged proteins were purified by nickel affinity chromatography. For bacterial expressions, pellets were resuspended in 30 mL 1× Ni-NTA buffer (50 mM Tris-HCl pH 7.5 at 4 °C, 300 mM NaCl), supplemented with 0.1 mg/mL lysozyme, 1 mg/ml cOmplete mini EDTA-free protease inhibitor (Roche) and 1 mM PMSF. This sample was incubated for 30 min at 4 °C with end-over-end mixing. Sonication was then performed on ice with an Ultrasonic Processor equipped with a microtip (Cole-Parmer), for four rounds for 60 s on 50% duty-cycle, separated by a 60 s pause. This crude lysate was then centrifuged for 45 min at 35,000 g at 4 °C to remove cell debris. The pH of the clarified lysate was adjusted to 7.5 using 1 M Tris-HCl pH 8.0.

For mammalian expressions, 10% (v/v) 10× Ni-NTA buffer (500 mM Tris-HCl pH 7.5 at 4 °C, 3 M NaCl) was added to filtered cell supernatants and pH was adjusted to 7.5 using 1 M Tris-HCl pH 8.0. cOmplete His-Tag Purification Resin (Roche) was settled in an Econo-Pac Chromatography Column (Bio-Rad) and washed with 20 column volumes (CV) of milliQ water, then 20 CV of Ni-NTA buffer. Washed resin was then added to the clarified lysate or mammalian supernatant in conical tubes and incubated for 1 h with rolling. Resin was settled by centrifugation at 500 g for 5 min, and supernatant was discarded. The resin was then washed on the column with 3× 40 CV of Ni-NTA wash buffer (10 mM imidazole in Ni-NTA buffer). To elute, 1 CV elution buffer (200 mM imidazole in Ni-NTA buffer) was added and incubated for 15 min, then collected by gravity flow. This elution was repeated without incubation another 5-10 times. Elution fractions were analyzed by SDS–PAGE with Coomassie staining, pooled and dialyzed for 16 h against 5,000-fold excess phosphate-buffered saline (PBS: 137 mM NaCl, 2.7 mM KCl, 10 mM Na_2_HPO_4_, 1.8 mM KH_2_PO_4_, pH 7.4). Proteins were then concentrated using a Vivaspin centrifugal concentrator (GE Healthcare) with appropriate molecular weight cut-off and stored at -80 °C.

Typical yields per L of mammalian cell culture were 3 mg for αCD22-SpyCatcher003, 4 mg for αCD22-RBD, 16-42 mg for RBD, 22 mg for RBD(L3), 6-22 mg for Spike, 13 mg for NTD, 25 mg for S2, 12 mg for mCD22 and 52 mg for FcγRIIb extracellular region.

### SpySwitch purification

Fabs bearing a SpyTag003 but no His-tag were purified at 4 °C by SpySwitch purification (Vester *et al*, 2022). Filtered cell supernatants were supplemented with 10% (v/v) 10× SpySwitch buffer (500 mM Tris-HCl, 3 M NaCl, pH 7.5 at 4 °C) and pH was adjusted to 7.5 using 1 M Tris-HCl pH 8.0. SpySwitch resin (0.75 mL per 30 mL cell supernatant) was equilibrated with 2×10 CV SpySwitch buffer (50 mM Tris-HCl pH 7.5, 300 mM NaCl), then incubated with filtered cell supernatant using an end-over-end rotator for 1 h at 4 °C. The resin was then washed with 4 × 10 CV SpySwitch buffer. pH-dependent elution was performed by incubating the resin with 6 × 1.5 CV SpySwitch elution buffer (50 mM acetic acid, 150 mM NaCl, pH 5.0) for 5 min each. The flow-through from each elution was immediately mixed with 6 × 0.3 CV of 1 M Tris-HCl pH 8.0 to neutralize the elution buffer. Proteins were dialyzed for 16 h against 5,000-fold excess PBS, then concentrated using a Vivaspin centrifugal concentrator (GE Healthcare) with molecular weight cut-off 20 kDa and stored at -80 °C. Typical yields per L of culture were 20-34 mg for the three anti-FcγRIIb Fabs, and 1.8-3.5 mg for anti-CD22-SpyTag003.

### Polyacrylamide gel electrophoresis

SDS-PAGE was performed using 8-16% polyacrylamide gels poured in 1.5 mm cassettes (Invitrogen). Prior to loading, protein samples were mixed with 6× loading buffer (0.23 M Tris HCl pH 6.8, 24% (v/v) glycerol, 120 μM bromophenol blue, 0.23 M sodium dodecyl sulfate). In addition, for reducing conditions, dithiothreitol (DTT) was added to a final concentration of 60 mM. Electrophoresis was performed at 180-200 V in 25 mM Tris-HCl pH 8.5, 192 mM glycine, 0.1% (w/v) SDS. PageRuler prestained (Thermo Scientific) protein ladder was used. Brilliant Blue G-250 was used to stain gels, which were then destained with MilliQ H_2_O. Gels were imaged on an iBright FL1500 Imaging System (Thermo Fisher) and analyzed using the iBright Analysis Software Versions 5.0.1 or 5.2.0 (Thermo Fisher). Mammalian-expressed proteins often showed a broad band relating to variable charge from multiple N-linked glycosylation sites.

### PNGase F deglycosylation

For deglycosylation of protein samples for SDS-PAGE, 4-6 µg glycoprotein was mixed with NEB Glycoprotein Denaturing Buffer (final concentration 0.5% (w/v) SDS and 40 mM DTT) in 10 µL total. This sample was incubated for 10 min at 100 °C, then briefly cooled on ice. Next, 2 µL NEB GlycoBuffer2 (final concentration 50 mM sodium phosphate, pH 7.5), 2 µL NP-40 (final concentration 0.5% (v/v)), and 6 µL MilliQ water were added to make up to 20 µL. Finally, 1 µL of 1 mg/mL PNGase F, made in-house recombinantly as previously described (Vankayala *et al*, 2026), was added and incubated for 1 h at 37 °C.

### VLP assembly

10-20 μM SpyCatcher003-mi3 was mixed at 1:0.95 molar ratio with SpyTagged proteins. For VLPs with two SpyTagged antigens, the antigens were pre-mixed in equal molar ratio. For inhibitory VLPs, Fab and antigen were pre-mixed in a 9:1 molar ratio. The reaction was left for 16 h in PBS on ice.

### DLS

VLP samples were centrifuged at 17,000 g for 30 min at 4 °C and transferred to a new tube. Samples were then diluted to 1-5 µM in PBS in a disposable cuvette (Malvern, ZEN0040). DLS was measured at 20 °C using a Zetasizer nano (Malvern). Before collecting data, the cuvette was incubated in the instrument for 2 min, to allow the sample temperature to stabilize. For each sample, three measurements were made, with the attenuation and number of runs per measurement automatically determined. Data were collected and analyzed in the Zetasizer Software v.7.13.

### Inhibitory bispecific assembly

Assembly was carried out in PBS (Driscoll *et al*, 2024). For DoubleCatcher-based inhibitory bispecifics, DoubleCatcher with molar concentration *N* (typically 50-60 µM) was incubated with 1.25 × *N* Fab-SpyTag003 for 3 h at 37 °C with shaking at 600 rpm. Then 3 × *N* Antigen-SpyTag003 and 1/12 × *N* MBP-superTEV were added to the reaction and incubated for 4 h at 34 °C with shaking at 600 rpm. SpyCatcher003 S49C resin (Driscoll *et al*, 2024) and amylose resin (New England Biolabs) were washed thrice in PBS and added to the reaction tube. This mixture was incubated for 16 h at 4 °C with end-to-end mixing to remove unreacted SpyTag003-linked proteins and MBP-superTEV from solution. Resin was removed by passing the sample through a filter column (Bio-Rad). For the DoubleCatchers shown in Fig. 5, GST-TEV protease was used instead of MBP-superTEV. GST-TEV was removed with glutathione-agarose (Sigma-Aldrich) instead of amylose resin. For Fab-SpyCatcher003 assemblies, Fab-SpyCatcher003 was mixed with excess SpyTag003-linked antigen and incubated for 4 h at 34 °C. Next, the reaction mixture was incubated with SpyCatcher003 S49C immobilized on SulfoLink resin (Thermo Fisher) for 16 h at 4 °C, with end-to-end mixing to remove excess unreacted SpyTag003-linked protein.

### Mass Spectrometry

MS was acquired using an Acquity UPLC System coupled to a Waters Corporation Xevo G2-S Q-TOF mass spectrometer. Protein samples were diluted to 1 µM in 30 μL PBS in UPLC autosampler vials (Fisher Scientific). Chromatography was performed using the Acquity UPLC Protein BEHC4 column (2.1 mm × 50 mm). A flow-rate of 0.2 mL/min was used and samples were eluted to the Q-TOF detector in deionized water with 0.1% (v/v) formic acid and 5% (v/v) acetonitrile. Each sample was analyzed once. Data were analyzed in the MassLynx software V4.2 SCN 971 (Waters). Waters’ maximum entropy deconvolution algorithm (MaxEnt 1) was used to process protein ionization data. Expected molecular weights for DoubleCatcher and GFP were calculated using the ExPASy ProtParam tool, with the N-terminal fMet removed. For GFP, fluorescent protein maturation was accounted for (-20 Da including loss of water and Tyr oxidation) (Tsien, 1998). Following purification of Fab-SpyTag003 constructs, we observed cleavage of the C-terminal lysine from SpyTag003 (-128 Da) and the conversion of N-terminal glutamine to pyroglutamate (-18 Da each) (Gadgil *et al*, 2006). For DoubleCatcher-assembled bispecific molecules, the loss of two water molecules (-36 Da) in the formation of two isopeptide bonds was factored in (Driscoll *et al*, 2024).

### Endotoxin depletion and quantification

All samples used in mouse immunizations were endotoxin-depleted. 1% (v/v) Triton X-114 was added to samples and mixed well. Tubes were incubated on ice for 5 min, then at 37 °C for 5 min, followed by 1 min centrifugation at 17,000 g to separate the phases. The aqueous phase supernatant was transferred to a new tube. This procedure was repeated 2 more times. A final round without Triton X-114 was performed to ensure that minimal Triton X-114 remained in the sample. Endotoxin quantification was performed with a Pierce Chromogenic Endotoxin Quant Kit (Thermo Fisher), according to manufacturer instructions. Final endotoxin concentrations were all below the endotoxin level threshold for vaccine products of 20 endotoxin units per mL (Brito & Singh, 2011; Hills *et al*, 2024).

### Mouse immunization and blood sampling

Animal experiments were performed according to the UK Animals (Scientific Procedures) Act 1986, under Project License PP9362617 granted to Tiong Kit Tan and approved by the University of Oxford Local Animal Welfare and Ethical Review Body. C57BL/6 female mice were obtained from Envigo and were 6 weeks old at the time of the first immunization. Mice were housed in accordance with the UK Home Office ethical and welfare guidelines and fed on standard chow and water ad libitum. Antigenic doses on VLPs were normalized to be molar equivalent to 0.5 µg RBD i.e. 17 pmoles per dose. Where two antigens were presented, both were at 17 pmoles. Spike dose was also 17 pmoles (∼ 2.4 µg). For inhibitory components, the undesired antigen was administered at a dose 40× in molar terms compared to that in the activator portion, except specified groups in Fig. 4, which received only 20× inhibitory components. Immunizations were performed intramuscularly in the gastrocnemius muscle on isoflurane (Abbott)-anesthetized mice.

For temporal separation experiments presented, prime inhibitory components were given at week 0, followed by the activatory GFP/RBD VLP at week 1, both at the same injection site. This was repeated at weeks 4 (inhibitory) and 5 (activatory) for the boost dose. For spatial-separation experiments (Fig. 3), the activatory GFP/RBD VLP was administered to the right gastrocnemius muscle and inhibitory components were administered to the left gastrocnemius muscle, on the same day at week 1.

For co-administration experiments, activatory and inhibitory portions of immunogens were mixed in PBS up to 25 µL per dose and these complete immunogens were mixed 1:1 (25 µL+ 25 µL) with either PBS (no adjuvant conditions) or AddaVax (InvivoGen), prior to administration. Boost was administered 4 weeks after prime.

Post-prime blood samples were obtained on week 4, the day before boost administration, through the tail vein using Microvette (CB300, Sarstedt) capillary tubes. Post-boost samples were obtained 8 weeks after initial immunization via cardiac puncture of humanely killed mice. The collected whole blood in microtainer SST tubes (Becton Dickinson) was allowed to clot at 23 °C for 30-60 min, before spinning down at 10,000 g for 6 min at 23 °C and storing at -20 °C. The sera were heat-inactivated at 56 °C for 30 min prior to analysis.

### Mouse antisera ELISA

Nunc MaxiSorp plates (Thermo Fisher) were coated with 50 µL per well of 80 nM purified antigen in PBS at 4 °C for 16 h. Plates were washed three times with PBST (1% v/v Tween 20 in PBS). Plates were blocked for 2-4 h in blocking buffer (5% (w/v) skimmed milk in PBS) at 25 °C. Plates were then washed three times with PBST. A serial dilution of sera was prepared by first diluting sera 1:100 into blocking buffer, then a series of seven 1:4 dilutions. For sera from mice immunized with Spike, the dilution series started at 1:50 rather than 1:100. Each sample:antigen combination was analyzed as a technical duplicate. Plates were incubated with sera for 1 h at 25 °C, then washed three times with PBST. Plates were incubated at 25 °C for 1 h with a 1:1,600 dilution of horseradish peroxidase-conjugated goat anti-mouse IgG antibody (Sigma-Aldrich, A4416). Plates were washed three times with PBST. Plates were then incubated at 25 °C for 5 min with 50 μL 1-Step Ultra TMB-ELISA Substrate Solution (Thermo Scientific). Next, 50 μL 1 M HCl was added to quench the reaction. Absorbance at 450 nm (A_450_) was measured by a FLUOstar Omega plate reader (BMG Labtech) using Omega MARS software (BMG Labtech).

A sigmoidal dose–response curve was fit to the absorbance data as previously described (Hills *et al*, 2024). Area under the curve was chosen because it better accounts for data across the range of values than endpoint titers (Hartman *et al*, 2018; Chivers *et al*, 2010). Results were visualized in GraphPad Prism (GraphPad Software v. 10.5.0).

### AviTag Biotinylation of recombinant receptors

BirA biotinylation was performed following published protocols (Fairhead & Howarth, 2015). Briefly, AviTag-linked protein (CD22 or FcγRIIb) at concentration C was mixed with C/15 molar concentration of GST-BirA, in the presence of 15C molar excess of D-Biotin, 5 mM MgCl_2_ and 1 mM ATP in PBS. This sample was incubated at 25 °C for 1 h, then the same amount of GST-BirA was added again and incubated for another hour. GST-BirA was then removed by incubating the reaction mixture with glutathione-agarose (Sigma-Aldrich) for 1 h at 4 °C under rotation. The now biotinylated protein was dialyzed for 16 h against 5,000-fold excess PBS to remove excess biotin. Quantitative biotinylation was confirmed by Traptavidin gel-shift on SDS-PAGE with Coomassie staining (Fairhead & Howarth, 2015).

### Receptor-binding ELISA

Nunc MaxiSorp plates (Thermo Fisher) were coated with 50 µL per well of 80 nM SpyCatcher002-MBP in PBS at 4 °C for 16 h. Plates were washed three times with PBST and blocked for 2-4 h with blocking buffer at 25 °C. Plates were then washed three times with PBST. SpyTag003-linked Fabs were added at 80 nM in blocking buffer and incubated for 30 min at 25 °C. Plates were washed three times with PBST. Biotinylated CD22-AviTag or biotinylated FcγRIIb-AviTag was added at 5, 50 or 150 nM in blocking buffer and incubated for 1 h at 25 °C. Plates were washed three times with PBST. Plates were incubated at 25 °C for 1 h with a 1:5,000 dilution of horseradish peroxidase (HRP)-conjugated Streptavidin (Pierce High-Sensitivity Streptavidin-HRP). Plates were washed three times with PBST. Plates were then incubated at 25 °C for 5 min with 50 μL 1-Step Ultra TMB-ELISA Substrate Solution (Thermo Scientific). Next, 50 μL 1 M HCl was added to quench the reaction. Absorbance at 450 nm (A_450_) was measured by a FLUOstar Omega plate reader (BMG Labtech) using Omega MARS software (BMG Labtech). Data were plotted using Microsoft Excel. v.16.89.1.

### Biolayer Interferometry

Kinetic experiments were performed on an Octet-Red BLI system (ForteBio) at 37 °C with 1,000 rpm shaking in BLI buffer (PBS with 0.05% (v/v) Tween 20). Assays were set up in 96-well plates (Greiner 655209) with 250 μL per well. Streptavidin biosensors (Sartorius, 18-5019) were placed in BLI buffer at least 15 min before use, to pre-hydrate. Biotinylated ectodomain of CD22 was loaded at 1 µg/mL to a total response of 0.75-1 nm. Association phase was performed for 400 s with 200 nM, 50 nM or 12.5 nM Fab, then dissociation was carried out for 400 s. For FcγRIIb, loading was performed with 0.5 µg/mL to a total response of 0.3-0.4 nm. Association phase was performed for 150 s with 100 nM, 50 nM, 25 nM or 12.5 nM Fab, then dissociation was carried out for 200 s. This shorter timeframe was chosen to minimize non-ideal behavior (Sherer & Cho, 2025). Analysis was performed in Octet DataAnalysis v.11.0.0.4. Reference subtraction was applied, with BLI buffer only instead of Fab in reference wells. A global fit was applied to association and dissociation across the 3 or 4 concentrations, with a 1:1 fitting model. We report the k_on_, k_off_, K_d_ and standard deviations automatically generated from this fit in the Octet software.

### Bioinformatics and Graphics Visualization

Protein structures were visualized using PyMOL (Schrödinger) version 2.3.4. Bispecific protein illustrations were assembled from predictions made using AlphaFold 3, accessed through the AlphaFold server. Each construct was predicted individually, then assembled in PyMOL by aligning Catchers and Tags to PDB 4MLI (Li *et al*, 2014). To visualize maximum reach, dihedral angles for flexible linkers were then manually set to φ = -135°, ψ = 135° and ω = 180°. The structure of SARS-CoV-2 Spike was from PDB ID 6VSB (Wrapp *et al*, 2020). This structure was input to ConSurf (Landau *et al*, 2005), accessed via the Colab notebook, to calculate sequence conservation. Multiple Sequence Alignment (MSA) was constructed from 32 Spike Homologs and conservation was mapped onto chain A of a Spike structure (PDB: 6VSB). Illustrations of constructs were based on structures predicted by ColabFold and designed using PixelmatorPro v.3.6.14.

### Statistics and Reproducibility

No statistical method was used to predetermine sample size. Significance across groups for ELISAs in Fig. 2E was calculated using a two-way ANOVA with Fisher’s Least Significant Difference (LSD) in GraphPad Prism (v. 10.5.0). Fisher’s LSD was used since only three groups were being compared and the overall group factor F-test was significant. Significance between antigens or groups in other serum ELISA experiments was calculated using multiple paired t-tests in GraphPad Prism (v. 10.5.0). Significance in the RBD-GFP titer values was also calculated by t-test, comparing each test group to antigenic-only control group. Stars were assigned as: * p < 0.05, ** p < 0.01, *** p < 0.001, no stars = non-significant. Experiments were not randomized and investigators were not blinded to allocation during experiments or analysis.

## Data availability

Sequences of constructs are available in GenBank, as described in the section “Plasmids and Cloning”. Plasmids that are deposited in the Addgene repository are described in the section “Plasmids and Cloning” (https://www.addgene.org/Mark_Howarth/). Further information and requests for resources and reagents should be directed to and will be fulfilled by the lead contact, M.R.H.

