## Supplementary Figures for "Duality Vaccination through Nanodesign of Click-Ligated B-Cell Inhibitory Receptor Bridging"

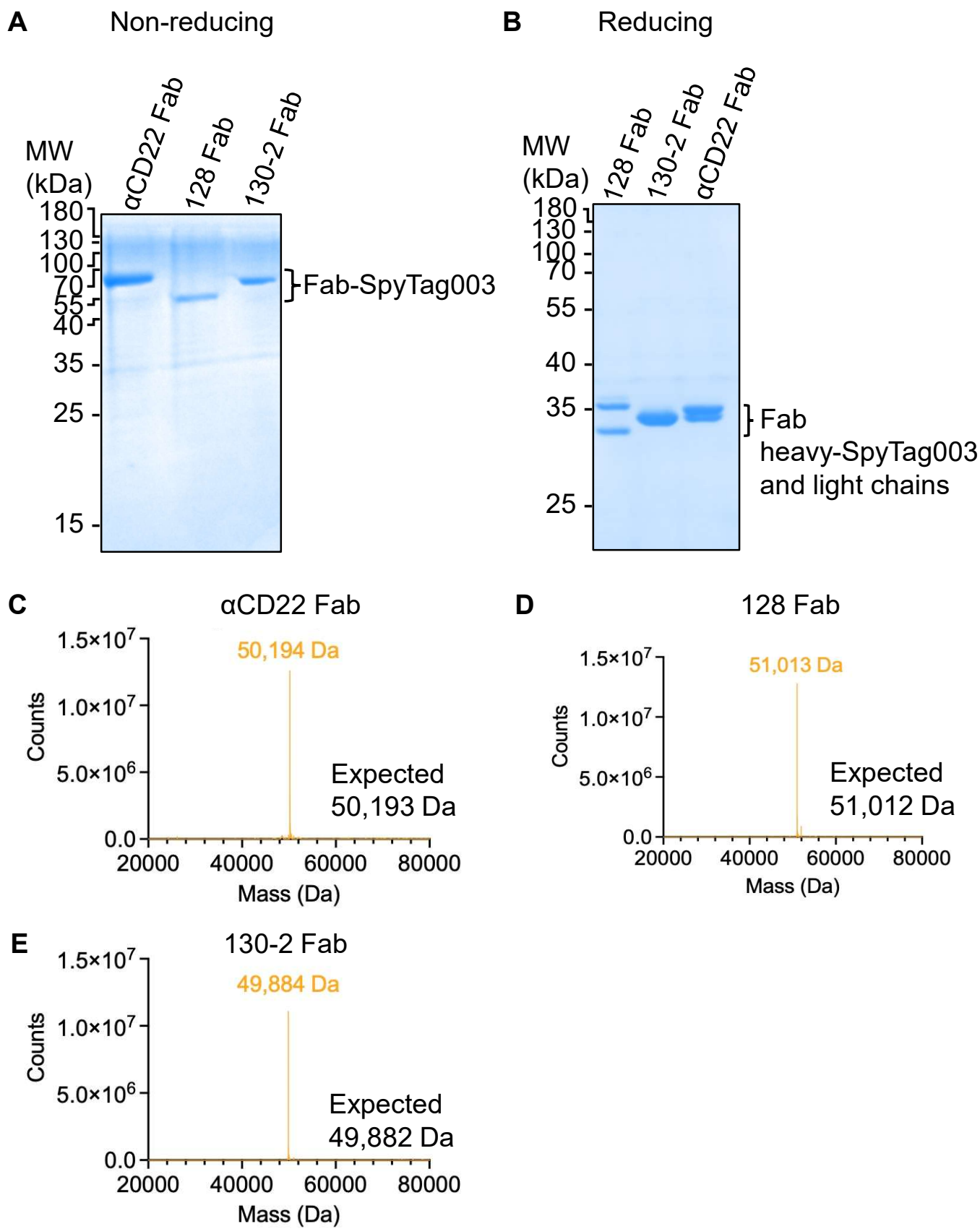

**Supplementary Fig. 1. Expression and purification of Fabs to inhibitory receptors.** (A) Purification of SpyTag003-linked Fabs. Fabs against CD22 or FcγRIIb (128 or 130-2) were purified from Expi293F cells and analyzed by non-reducing SDS-PAGE with Coomassie staining. (B) Reducing SDS-PAGE of Fabs as in (A). (C) Electrospray Ionization Mass Spectrometry of anti-CD22 Fab, with observed peak mass shown alongside the expected mass. (D) Mass as in (C) for anti-FcγRIIb Fab 128. (E) Mass as in (C) for anti-FcγRIIb Fab 130-2.

**A**

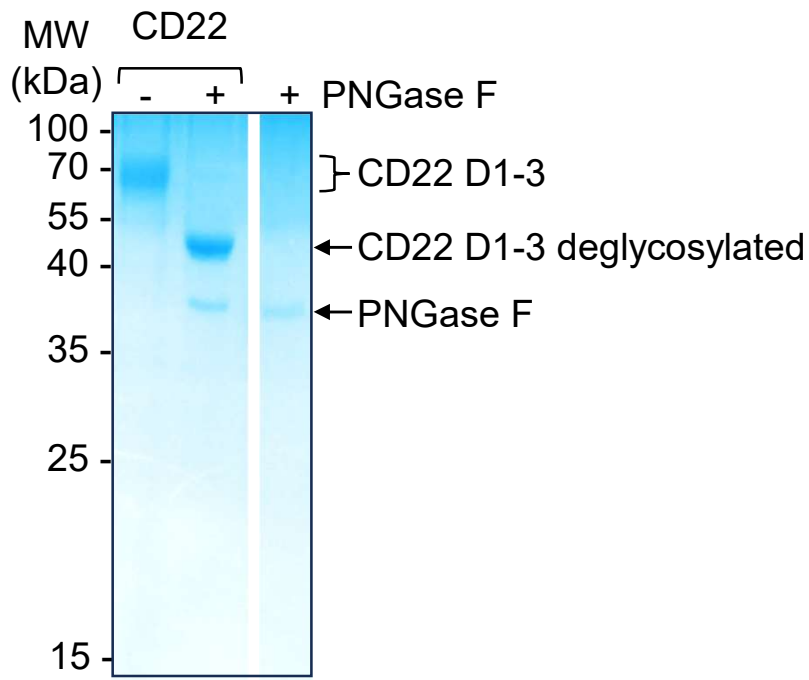

CD22 domain organization

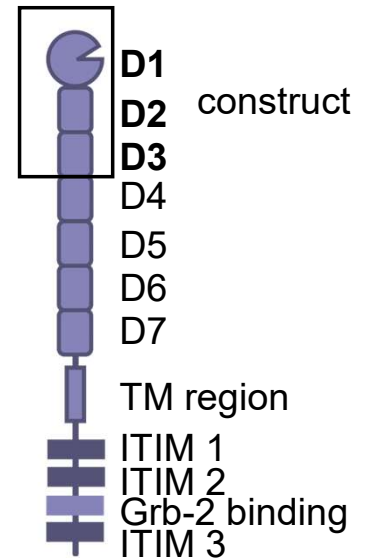

**B**

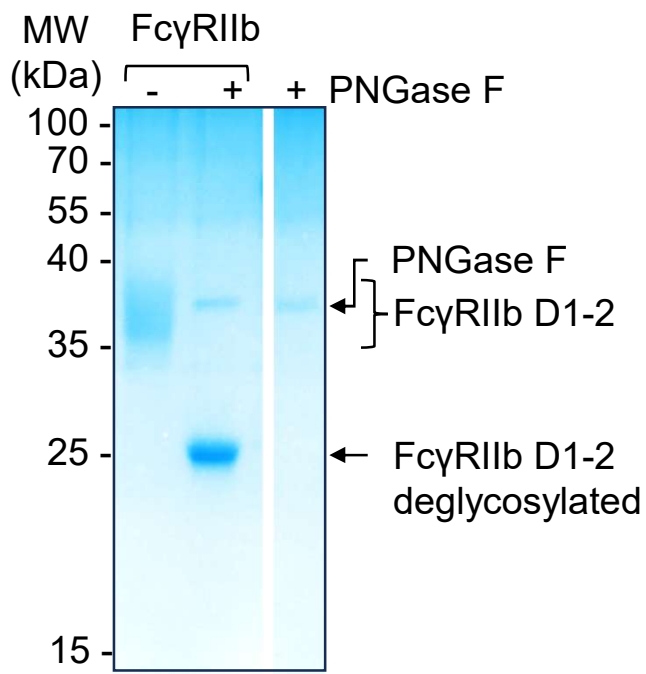

FcγRIIb domain organization

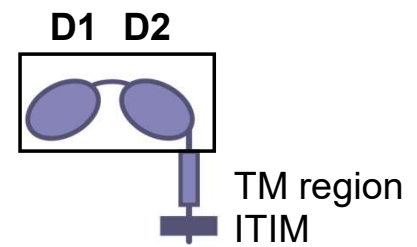

**Supplementary Fig. 2. Purification of inhibitory receptor ectodomains. (A)** CD22 extracellular domains 1-3 (D1-3) were cloned for expression in Expi293F cells and analyzed by SDS-PAGE with Coomassie staining. **(B)** FcγRIIb extracellular domains 1-2 (D1-2) were cloned for expression in Expi293F cells and analyzed by SDS-PAGE with Coomassie staining. In the lanes without deglycosylation by Peptide:N-glycosidase F (PNGase F), the broad band relates to variable charge from multiple N-linked glycosylation sites.

**A ELISA anti-CD22 Fab binding to CD22**

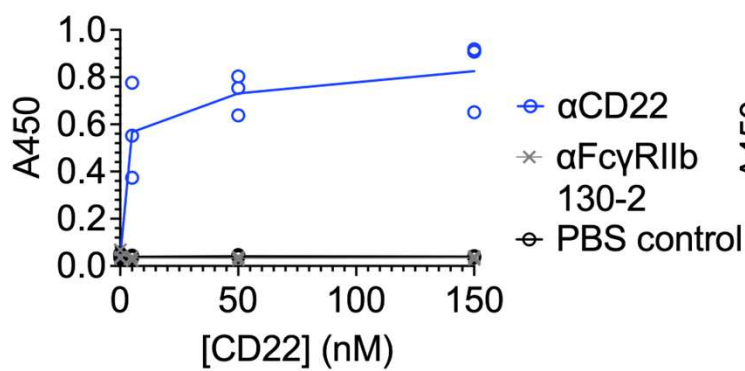

**B ELISA Fabs binding to FcγRIIb**

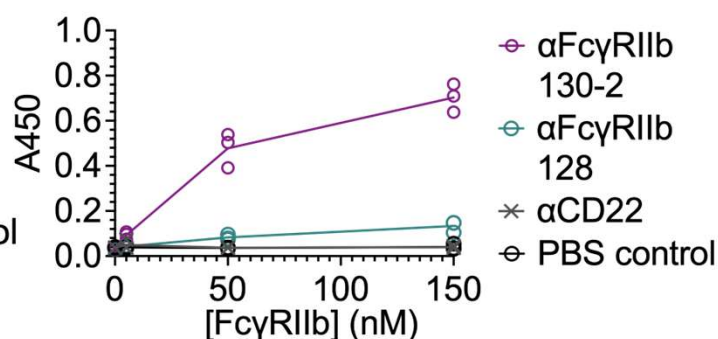

**C BLI binding anti-CD22 to CD22**

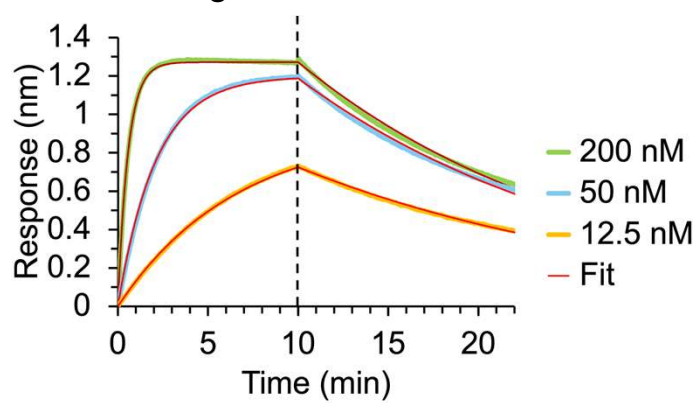

**D BLI binding 130-2 to FcγRIIb**

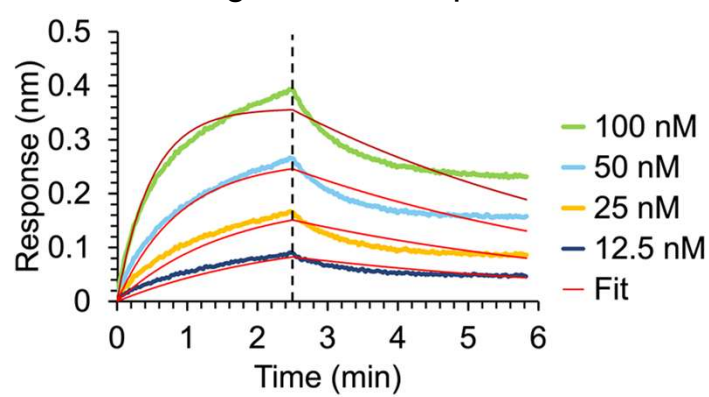

**E BLI binding 128 to FcγRIIb**

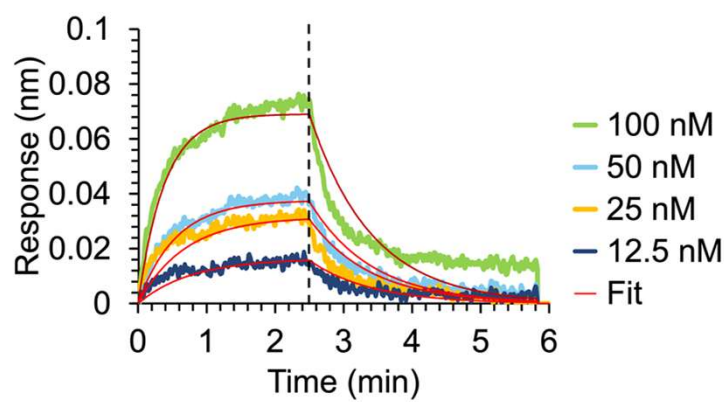

**F BLI summary**

| | $K_d$ (nM) | $k_{on}$ ($M^{-1}s^{-1}$) | $k_{off}$ ($s^{-1}$) |
| --- | --- | --- | --- |
| **α-CD22 to CD22** | $7.1 \pm 0.1$ | $1.4 \pm 0.1 \times 10^5$ | $9.9 \pm 0.1 \times 10^{-4}$ |
| **130-2 to FcγRIIb** | $10 \pm 0.3$ | $3.1 \pm 0.1 \times 10^5$ | $32 \pm 0.4 \times 10^{-4}$ |
| **128 to FcγRIIb** | $53 \pm 2.8$ | $3.6 \pm 0.2 \times 10^5$ | $190 \pm 2.8 \times 10^{-4}$ |

**Supplementary Fig. 3. Validation of Fab binding to inhibitory receptor ectodomains.** (A) ELISA showing anti-CD22 Fab binding to CD22 ectodomain. Anti-CD22 Fab or controls PBS or 130-2 Fab were immobilized on the plate, before varying concentrations of site-specifically biotinylated CD22 ectodomain were added. Detection was then performed colorimetrically using streptavidin-HRP. Each value was plotted for 3 replicate wells with the line connecting the mean. (B) ELISA showing anti-FcγRIIb Fab binding to FcγRIIb ectodomain, conducted as in (A). (C) BLI of anti-CD22 Fab binding to CD22 ectodomain. Fab was added at 12.5, 50 or 200 nM. Data are overlaid with the model fit in red. (D) BLI of 130-2 Fab binding to FcγRIIb ectodomain. Fab was added at 12.5, 25, 50 or 100 nM. Non-ideal binding behavior is likely caused by aggregation at the sensor surface. (E) BLI of 128 Fab binding to FcγRIIb ectodomain, as in (D). (F) Summary of BLI, showing $K_d$, $k_{on}$ and $k_{off}$ for Fab binding (mean ± 1 s.d.).

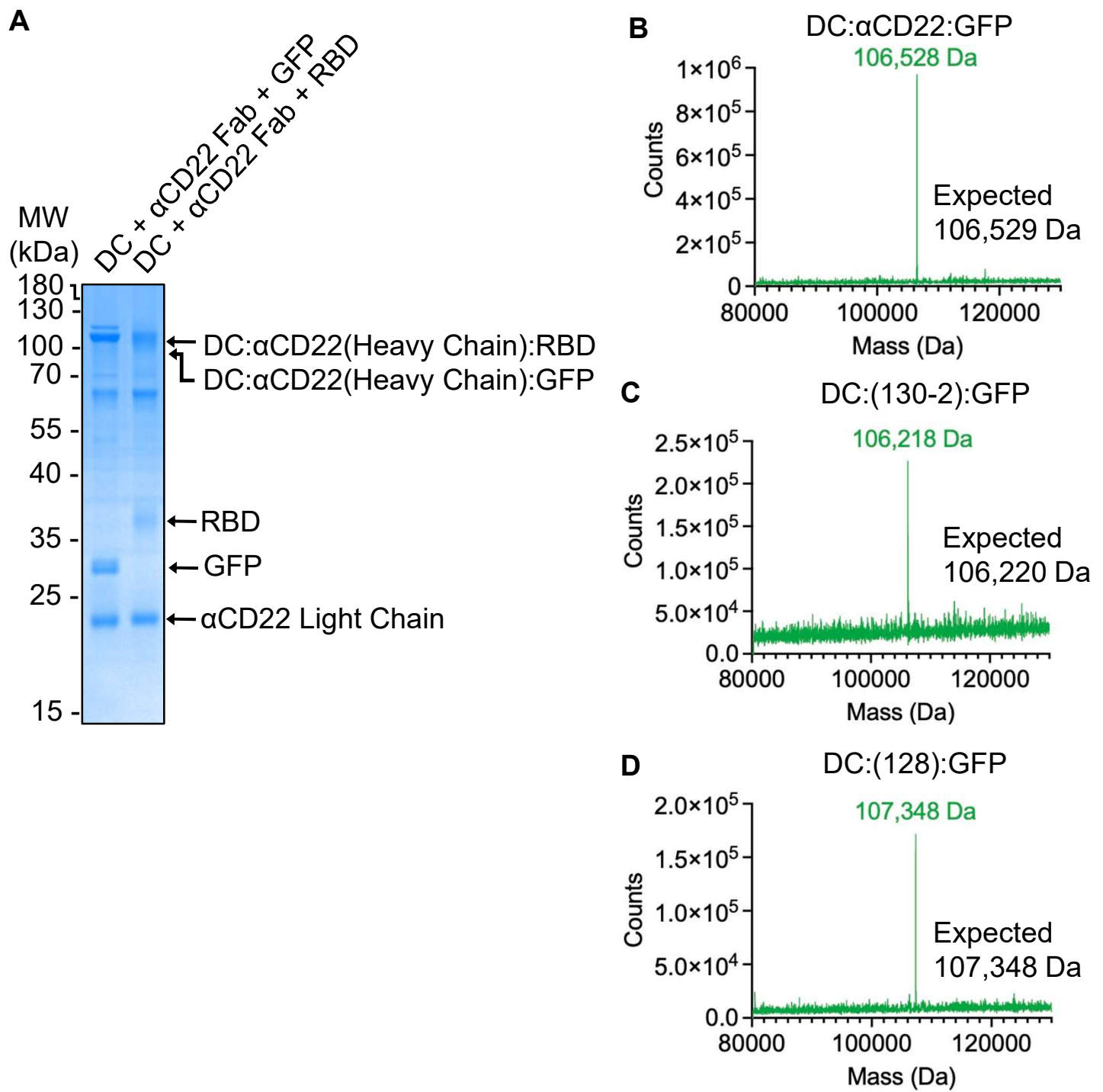

**Supplementary Fig. 4. Analysis of inhibitory immunogen assembly.** (A) Assembly of DoubleCatcher (DC) coupled to anti-CD22 Fab and RBD or GFP, analyzed by reducing SDS-PAGE with Coomassie staining. (B) Electrospray Ionization Mass Spectrometry of DoubleCatcher coupled to GFP and anti-CD22 Fab, with observed peak mass shown alongside the expected mass. (C) Mass as in (B) for DoubleCatcher coupled to GFP and anti-FcγRIIb Fab 130-2. (E) Mass as in (B) for DoubleCatcher coupled to GFP and anti-FcγRIIb Fab 128.

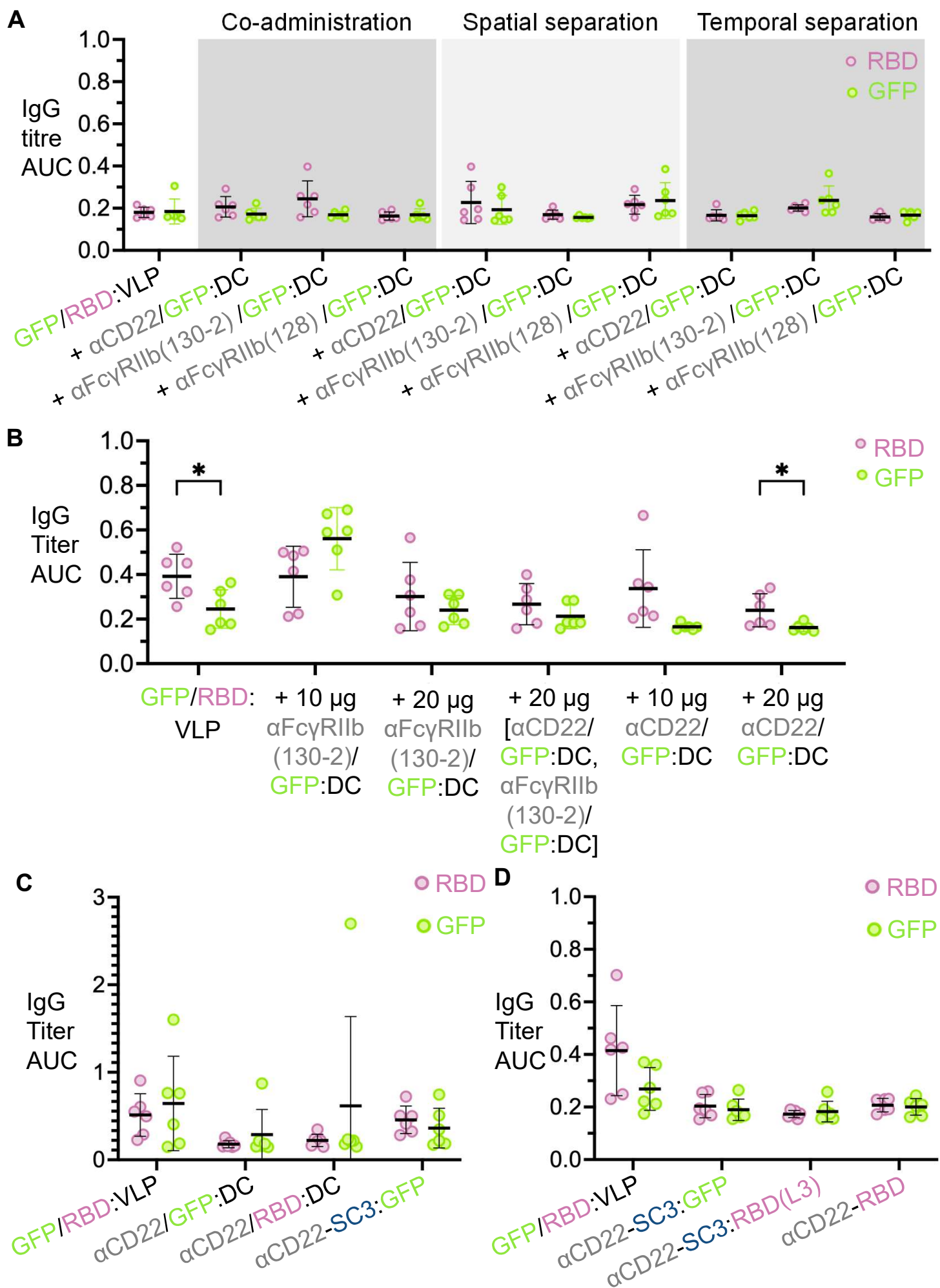

Supplementary Fig. 5 (start)

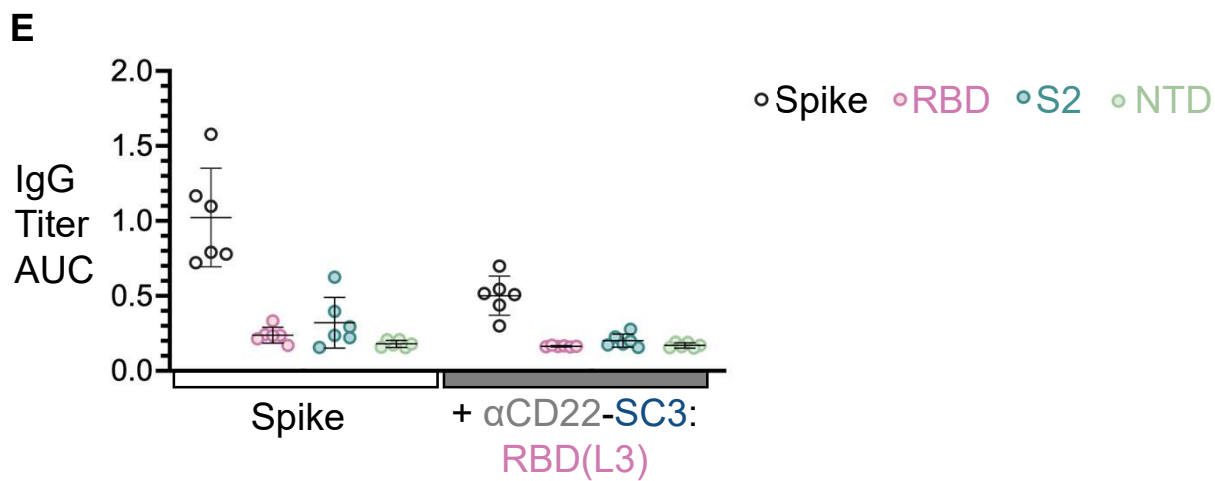

**Supplementary Fig. 5. Antibody responses following priming.** (A) Antibody responses following priming, in line with the boost in Fig. 3C. (B) Antibody responses following priming, in line with the boost in Fig. 4B. (C) Antibody responses following priming, in line with the boost in Fig. 5B. (D) Antibody responses following priming, in line with the boost in Fig. 5G. (E) Antibody responses following priming, in line with the boost in Fig. 6E.

**A**

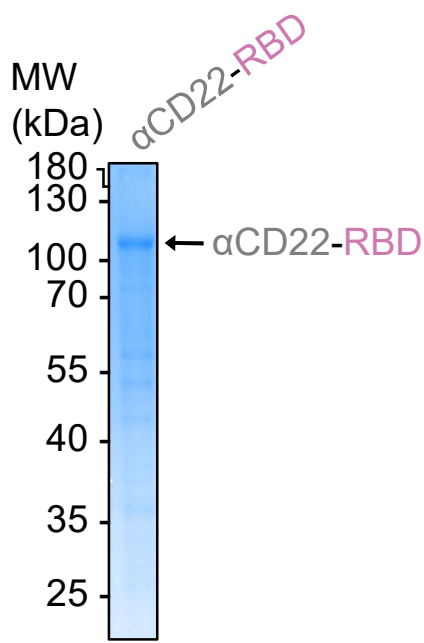

**B**

| Construct | Protein illustration | Total linker length |
| --- | --- | --- |
| DoubleCatcher:<br>αCD22:GFP |  | 26 aa |
| αCD22-SpyCatcher003:<br>GFP |  | 17 aa |
| DoubleCatcher:<br>αCD22:RBD |  | 54 aa |
| αCD22-SpyCatcher003:<br>RBD(L3) |  | 14 aa |
| αCD22-RBD |  | 3 aa |

**Supplementary Fig. 6. Modulating the size of inhibitory immunogens.** (A) Expression and purification of a direct genetic fusion of anti-CD22 Fab to RBD, analyzed by non-reducing SDS-PAGE with Coomassie staining. (B) Illustration of potential extended length of inhibitory immunogens, from AlphaFold prediction of individual units and modeling extended linker conformation. Total linker length is calculated from the Glycine-Serine spacers.

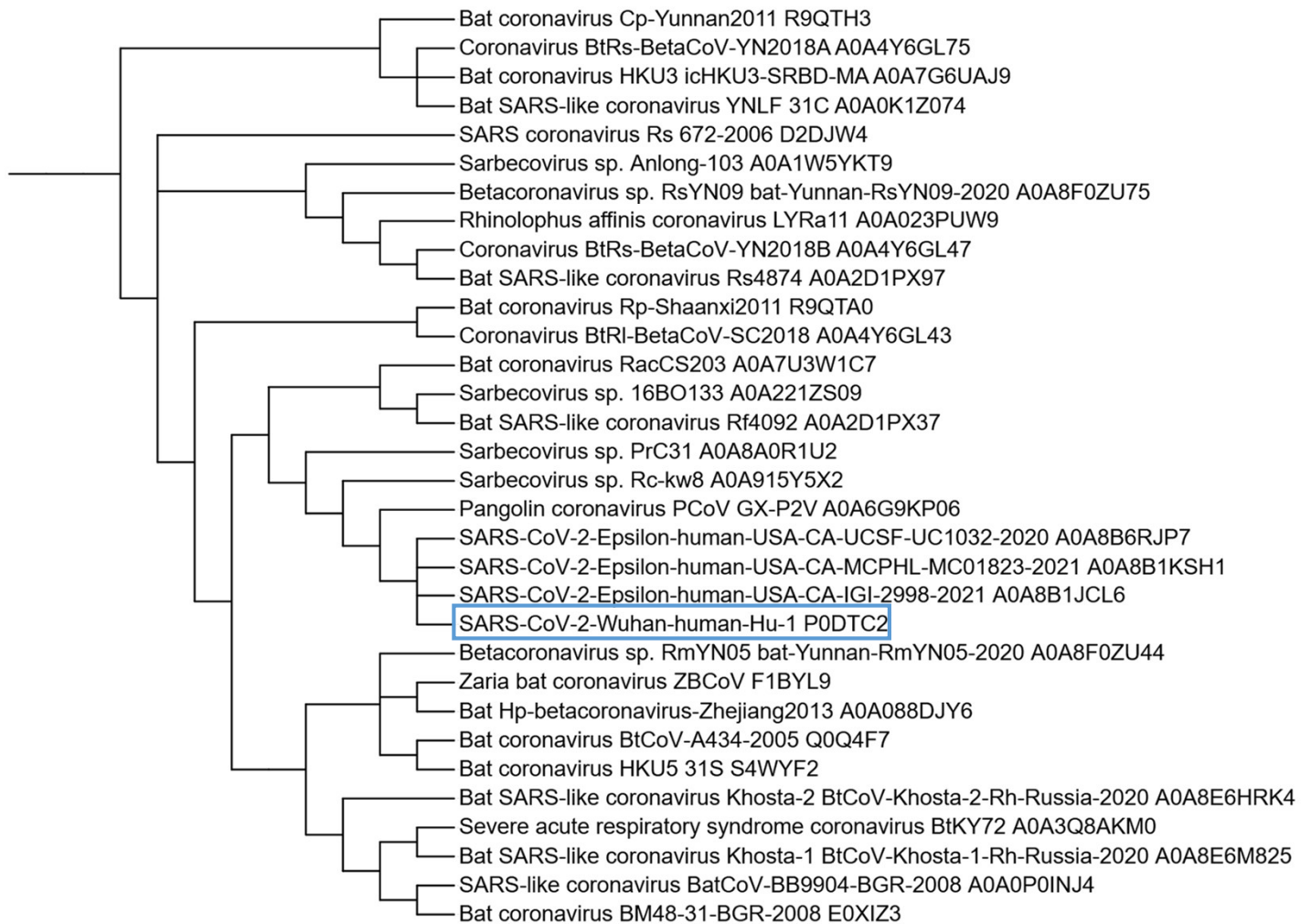

**Supplementary Fig. 7. Phylogenetic tree of sequences used in Spike conservation calculation.** Labels follow the format Virus\_strain-or-isolate, with the trailing code being the UniProt/UniParc accession number for that sequence.
